# Status of aquatic and riparian biodiversity in artificial lake ecosystems with and without management for recreational fisheries: implications for conservation

**DOI:** 10.1101/667493

**Authors:** Robert Nikolaus, Malwina Schafft, Andreas Maday, Thomas Klefoth, Christian Wolter, Robert Arlinghaus

**Affiliations:** Department of Biology and Ecology of Fishes, Leibniz-Institute of Freshwater Ecology and Inland Fisheries, Berlin, Germany; Angler Association of Lower Saxony, Hannover, Germany; Division for Integrative Fisheries Management, Albrecht Daniel Thaer-Institute of Agriculture and Horticulture, Faculty of Life Science, Humboldt-Universität zu Berlin, Germany

**Keywords:** amphibians, biodiversity, birds, disturbance, fishing, lake, littoral, recreation, riparian, vegetation

## Abstract

1. Humanity is facing a biodiversity crisis, with freshwater-associated biodiversity in a particularly dire state. Novel ecosystems created through human use of mineral resources, such as gravel pit lakes, can provide substitute habitats for conservation of freshwater and riparian biodiversity. However, many of these artificial ecosystems may exhibit high recreational use intensity, which may limit their biodiversity potential.
2. The species richness of several taxa (plants, amphibians, dragonflies, damselflies, waterfowl, songbirds) was assessed and a range of taxonomic biodiversity metrics were compared between gravel pit lakes managed for recreational fisheries (N = 16) and unmanaged reference lakes (N = 10), while controlling for non-fishing related environmental variation.
3. The average species richness of all examined taxa was similar among both lake types and no substantial differences in species composition were revealed when examining the pooled species inventory. Similarly, there were no differences among lake types in the presence of rare species and in the Simpson diversity index across all the taxa that were assessed.
4. Variation in species richness among lakes was correlated with woody habitat, lake morphology (surface area and steepness) and land use, but not correlated with the presence of recreational fisheries. Thus, non-fishing related environmental variables had stronger effects on local species presence than recreational-fisheries management or the presence of recreational anglers.
5. Collectively, no evidence was found that anglers and recreational-fisheries management constrain the development of aquatic and riparian biodiversity in gravel pit lakes in the study region. Conservation of species diversity at gravel pit lakes could benefit from an increasing reliance on habitat enhancement activities.

## 1. Introduction

Globally, biodiversity is in steep decline, with an estimated 1 million species currently threatened by extinction (Díaz et al., 2019). The biodiversity decline is particularly prevalent in freshwaters (Reid et al., 2019), where habitat alteration and fragmentation, pollution, biological invasions and climate change are key drivers (Dudgeon et al., 2006).

Artificially created aquatic habitats, such as gravel pit lakes or ponds, could maintain and increase native freshwater biodiversity by providing refuge and secondary habitats for rare or endangered species (Damnjanović et al., 2018; Oertli, 2018). The origins of artificial lake ecosystems are often relatively recent (less than 100 years of age; Zhao, Grenouillet, Pool, Tudesque, & Cucherousset, 2016), where artificial lakes are often created by mining for mineral resources (Saulnier-Talbot & Lavoie, 2018). More than one billion tonnes of sand and gravel were excavated in more than 24,500 quarries and pits within the European Union in 2017 alone (European Aggregates Association (UEPG), 2017). The resulting numerous artificial lakes (for simplicity henceforth referred to as gravel pit lakes) have become common elements in many cultural landscapes across the industrialized world (Oertli, 2018).

Lakes, including gravel pit lakes, provide many ecosystem services to humans. These include provisioning services, such as fish yield, as well as a range of cultural services, such as recreation (Meyerhoff, Klefoth, & Arlinghaus, 2019; Venohr et al., 2018). Although the benefits of water-based recreation can be substantial, water-based activities can also negatively impact the biodiversity of freshwater ecosystems (Venohr et al., 2018). For example, human activities can reduce littoral and riparian habitat quality and thereby negatively affect associated taxa (Spyra & Strzelec, 2019). Water-based recreation has also been found to negatively impact birds through fright responses to humans (Dear, Guay, Robinson, & Weston, 2015), dogs (Randler, 2006) or pleasure boats (McFadden, Herrera, & Navedo, 2017). Therefore, the management of gravel pit lakes and other artificial waterbodies would benefit from the joint consideration of the well-being aquatic recreation generates for humans and the possible negative biodiversity impacts from aquatic recreation.

Many gravel pit lakes located in central Europe are used for recreational fisheries (Matern et al., 2019; Zhao et al., 2016). In some regions of the world anglers are not only resource users, but also managers of fish populations and habitats (Arlinghaus, Müller, Rapp, & Wolter, 2017). This particularly applies to Germany, where organizations of anglers, usually angling clubs and associations, are leaseholders or owners of freshwater fishing rights, and in this position are also legally entitled to manage fish stocks (Arlinghaus, Müller, et al., 2017). This entails the sovereignty to stock fish, to manage littoral habitat and introduce access and harvest regulations (Arlinghaus, Müller, et al., 2017). Because stocking of fish is particularly prevalent in freshwater recreational-fisheries management, key impacts of the presence of recreational fisheries and associated management activities can be expected at the fish stock and fish community levels (Matern et al., 2019; Zhao et al., 2016). Angler-induced changes typically elevate fish species richness through the release and introduction of large-bodied “game” fishes of high fisheries interest (Matern et al., 2019; Zhao et al., 2016). In turn, the altered fish community may affect submerged macrophytes, (e.g., due to introduction of benthivorous fish that uproot macrophytes; Bajer et al., 2016), and other taxa through predation (e.g., birds: Cucherousset et al., 2012; amphibians: Hecnar & M’Closkey, 1997; or invertebrates: Knorp & Dorn, 2016). In addition, anglers may modify littoral habitats to create access to angling sites, thereby affecting species richness of plants (O’Toole, Hanson, & Cooke, 2009) and dragonflies (Z. Müller et al., 2003), or affecting mobile taxa, such as birds, through direct contact and disturbances (Bell, Delany, Millett, & Pollitt, 1997; Cryer, Linley, Ward, Stratford, & Randerson, 1987). Indirectly, angler presence can also inadvertently kill non-targeted wildlife, e.g., through lost fishing gear that is ingested by birds or where birds become entangled (Franson et al., 2003; Sears, 1988). Therefore, anglers can both be seen as stewards of aquatic ecosystems (Granek et al., 2008) as well as a potential threat to certain aquatic taxa depending on the local angling intensity and other conditions (Reichholf, 1988).

In Germany, fisheries (including recreational angling) are regulated by fisheries laws specific to the Federal state, while the protection of species and habitat types is regulated by Federal and state-specific nature conservation legislation. Conflicts with angler interests regularly occur when nature conservation authorities implement rules that partially or fully constrain access to water bodies to achieve conservation goals (Arlinghaus, 2005). Conservation-motivated constraints of angling or recreational-fisheries-management actions (e.g., stocking) are increasingly applied within artificial lake ecosystems through the implementation of national or international conservation law (e.g., the European Habitats Directive; Council of the European Communities, 1992). For example, in some regions of Germany recreational fisheries have been excluded from follow-up use of newly created gravel pit lakes during the process of licensing the sand or gravel extraction (H. Müller, 2012). Such bans of future angling use are often justified by the assumption that angling is particularly impactful for disturbance-sensitive taxa (e.g., waterfowl) or for habitats of special conservation concern (H. Müller, 2012; Reichholf, 1988).

To contribute to this ongoing debate, the work presented here has studied the taxonomic biodiversity associated with gravel pit lakes using a space-for-time substitution design comparing lakes managed and used by recreational fisheries with lakes that do not experience recreational fisheries actions and therefore lack angler impacts. The study goal was to examine the impact of recreational fisheries on the taxonomic aquatic and riparian biodiversity detectable at typical gravel pit lakes in north-western Germany. The specific objective was to estimate the effect of recreational fisheries on species richness, faunal and floral composition, community diversity and conservation value across a range of aquatic and riparian taxa that are protected by national and European conservation legislation (e.g., birds, amphibians, dragonflies). Because the absence of recreational fisheries in a given gravel pit lake does not mean the ecosystems remain undisturbed from other recreational uses (e.g., swimming, walking), it was hypothesized that the presence of recreational fisheries and associated management activities would, on average, not affect species richness and conservation value of taxa that are not specifically targeted by anglers (Odonata, amphibians, submerged and riparian vegetation, waterfowl and songbirds). This hypothesis was formulated as a statistical null hypothesis to be falsified by empirical data.

## 2. Methods

### Study area and lake selection

This study was conducted in the Central Plain ecoregion of Lower Saxony in north-western Germany (Figure 1), where natural lentic waters are scarce. Of 35,048 ha of total standing waters in Lower Saxony 73 % by area and more than 99 % by number are artificial lakes. These artificial waterbodies consist of mainly ponds and small gravel pit lakes with less than 10 ha surface area (Nikolaus et al., in review).

**Figure 1:**
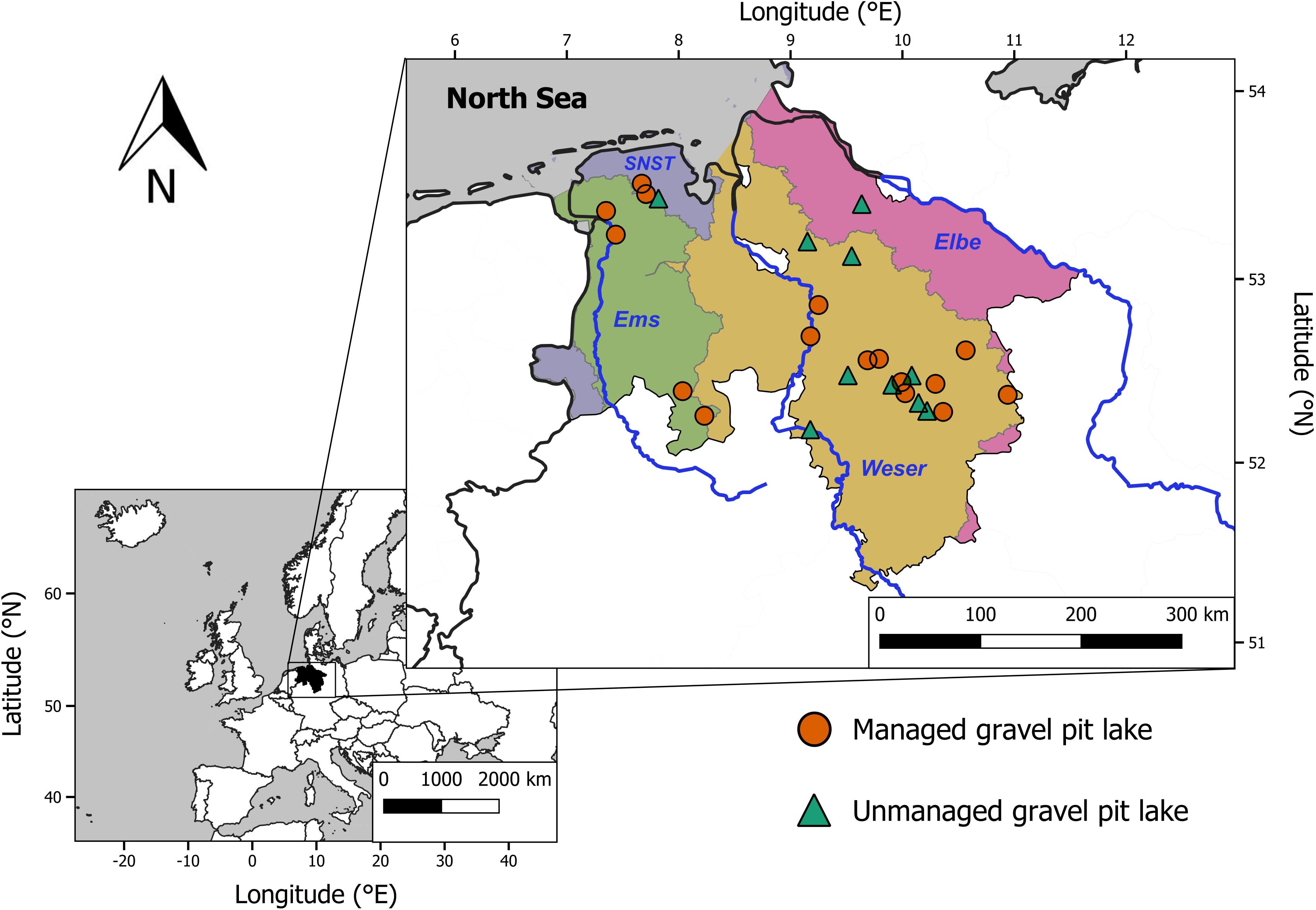
Map of study sites in Lower Saxony (Germany) together with the catchments (green = Ems, orange = Weser, magenta = Elbe, blue = SNST/small North Sea tributaries) and main rivers (dark blue; Ems, Weser, Elbe).

Most gravel pit lakes in Lower Saxony, and Germany as a whole, are managed for recreational fisheries by angler associations and clubs. These lakes are thus exposed to regular stocking with species of fisheries interest and are subject to access and harvest rules, regular controls by fisheries inspectors, and fishing club activities like collecting litter and cleaning and development of the littoral zone (Arlinghaus, Müller, et al., 2017). Similar activities are largely absent in gravel pit lakes not used for recreational fisheries, which are much rarer in number but still occur in Lower Saxony and elsewhere across Germany. For this study, a set of gravel pit lakes managed by recreational fisheries (defined as managed lakes) was selected, and compared to another set of gravel pit lakes not experiencing any form of legal angling and recreational fishing-related management (defined as unmanaged lakes, Table 1).

**Table 1:** Descriptors of gravel pits sampled in Lower Saxony. The locations of the catchments (fourth column) are shown in Figure 1. Trophic state was determined using Riedmüller, Hoehn, & Mischke (2013). The pH values and conductivities are annual means.

| No. | Lake name | Lake type | catchment | End of mining | Trophic state | Maximum depth [m] | Lake area [ha] | pH value | Conductivity [$\mu\text{S}\cdot\text{cm}^{-1}$] | Main recreationists and activities identified during on-site visits |
| --- | --- | --- | --- | --- | --- | --- | --- | --- | --- | --- |
| 01 | Chodhemster Kolk | Managed | Ems | 1971 | Mesotrophic | 10.1 | 3.2 | 6.7 | 208 | dog walkers |
| 02 | Collrunge | Managed | SNST<sup>1</sup> | 1982 | Mesotrophic | 8.6 | 4.3 | 7.9 | 200 | Anglers, dog walkers |
| 03 | Donner Kiesgrube 3 | Managed | Weser | 2000 | Eutrophic | 5.2 | 1.0 | 7.8 | 632 | Cyclists |
| 04 | Kiesteich Brelingen | Managed | Weser | 1999 | Mesotrophic | 8.7 | 8.5 | 7.5 | 299 | Anglers, dog walkers |
| 05 | Kolshorner Teich | Managed | Weser | 1980 | Mesotrophic | 16.1 | 4.3 | 7.6 | 547 | Anglers, horse riders |
| 06 | Linner See | Managed | Ems | 2000 | Mesotrophic | 11.2 | 17.7 | 8.5 | 301 | Anglers |
| 07 | Meitzer See | Managed | Weser | 2006 | Oligotrophic | 23.5 | 19.5 | 7.9 | 618 | Anglers, dog walkers, swimmers |
| 08 | Neumanns Kuhle | Managed | Ems | 1970 | Polytrophic | 6.2 | 6.9 | 8.6 | 585 | Anglers, dog walkers |
| 09 | Plockhorst | Managed | Weser | 1998 | Eutrophic | 8.2 | 14.3 | 8.0 | 338 | Anglers, dog walkers, horse riders |
| 10 | Saalsdorf | Managed | Weser | 1995 | Mesotrophic | 9.2 | 9.0 | 9.0 | 566 | Anglers, dog walkers |
| 11 | Schleptruper See | Managed | Ems | 1965 | Mesotrophic | 10.1 | 4.0 | 8.2 | 502 | Anglers, dog walkers |
| 12 | Stedorfer Baggersee | Managed | Weser | 1983 | Eutrophic | 2.8 | 1.9 | 7.5 | 352 | Anglers, dog walkers |
| 13 | Steinwedeler Teich | Managed | Weser | 1978 | Mesotrophic | 9.1 | 10.4 | 7.5 | 742 | Anglers, dog walkers, cyclists |
| 14 | Wahle | Managed | Weser | 1990 | Mesotrophic | 12.1 | 8.1 | 7.6 | 728 | Anglers, dog walkers, cyclists, horse riders |
| 15 | Weidekampsee | Managed | Weser | 1994 | Oligotrophic | 4.3 | 2.9 | 8.0 | 461 | Anglers, dog walkers |
| 16 | Wiesedermeer | Managed | SNST<sup>1</sup> | 1990 | Mesotrophic | 9.2 | 2.9 | 8.0 | 136 | Anglers, dog walkers |
| 17 | Bülstedt | Unmanaged | Weser | 1991 | Polytrophic | 1.1 | 2.4 | 7.6 | 285 | Birdwatchers |
| 18 | Goldbeck | Unmanaged | Elbe | 1992 | Eutrophic | 5.0 | 2.3 | 7.9 | 293 | Owner/friends, horse riders, swimmers |
| 19 | Handorf | Unmanaged | Weser | 2004 | Eutrophic | 23.0 | 13.6 | 9.2 | 666 | Horse riders, dog walkers |
| 20 | Hänigsen | Unmanaged | Weser | 2011 | Mesotrophic | 12.3 | 6.2 | 8.6 | 437 | Swimmers, dog walkers, anglers, campfires, dumping waste |
| 21 | Heeßel | Unmanaged | Weser | 1963 | Eutrophic | 7.4 | 0.9 | 8.2 | 537 | No |
| 22 | Hopels | Unmanaged | SNST<sup>1</sup> | 1998 | Mesotrophic | 14.5 | 5.5 | 7.7 | 215 | Swimmers, dog walkers |
| 23 | Lohmoor | Unmanaged | Weser | 1991 | Eutrophic | 7.4 | 4.1 | 8.2 | 159 | Birdwatchers |
| 24 | Pfütze | Unmanaged | Weser | 2000 | Mesotrophic | 7.3 | 10.6 | 7.7 | 342 | Dog walkers, canoeing, swimmers, cyclists |
| 25 | Schwicheldt | Unmanaged | Weser | 2007 | Mesotrophic | 10.0 | 1.7 | 8.4 | 895 | Owner/hunter, dog walkers |
| 26 | Xella | Unmanaged | Weser | 1975 | Mesotrophic | 7.3 | 2.1 | 7.5 | 957 | No (restricted access) |

<sup>1</sup> Small North Sea Tributaries

Managed lakes were identified through a survey of all angling clubs organized in the Angler Association of Lower Saxony. Lakes were selected according to the following criteria: the lake was owned by a fishing club, of small size (1-20 ha) and had no dredging in the last ten years (“old age”). This approach yielded N = 16 managed lakes as study sites spread across Lower Saxony in 10 angling clubs (Table 1, Figure 1).The angler density (number of anglers per unit water area) for these clubs ranged from 8 to 43 anglers per hectare (mean ± SE: 21 ± 3.6 anglers per ha). These angler densities correspond to averages known for German and Lower Saxonian angling clubs, which are 24 ± 2.5 and 22 ± 10.8 anglers per ha, respectively. All selected angler-managed lakes experienced regular angling activities and fisheries-management actions, including annual stocking of a range of fish species and regular shoreline development activities, such as mowing of angling sites and litter removal.

Gravel pits not managed by and for anglers were identified in close vicinity to the managed lakes where available (Figure 1). The number of unmanaged lakes in the state was much smaller than the number of managed lakes. Overall, N = 10 unmanaged lakes were identified, which were of similar age, size and other environmental conditions to the managed lakes, but differed from the managed ones by the absence of an angling club and any form of legal angling and fisheries-management for at least five years prior to the onset of our study (Table 1, Figure 1). Both lake types were accessible to non-angling recreation as they were not fenced.

In a subset of the selected lakes, Matern et al. (2019) previously conducted fish faunistic surveys revealing identical fish abundances and biomasses in both lakes types, but larger local fish species richness and significantly more abundant game fishes (particularly predators and large-bodied cyprinids such as carp, Cyprinus carpio) in managed lakes compared to unmanaged ones. These data showed that the angler-managed lakes included in this study were indeed more intensively managed in terms of fish stocking and hosted a substantially different fish community. This finding was a relevant precondition of the study design in that managed and unmanaged lakes differed in traces left by fisheries-management and fisheries use, both in terms of fish community composition and angler presence in the littoral zone.

Despite the attempt to select lakes as similar as possible in terms of the environment (e.g., age, surface area, trophic state), a set of environmental variables was assessed and integrated into the statistical analyses to more carefully isolate the possible impact of recreational-fisheries management on biodiversity, while controlling for other key environmental differences among lakes that could also affect the community composition of specific taxa (e.g., morphometry, land use, habitat structure).

### Land use

Several indicators of land use and spatial arrangement in Lower Saxony across catchments were assessed. Shortest-path distances of lakes to nearby cities, villages, lakes, canals and rivers were calculated in Google Maps (© 2017). Subsequently, a share of different land use categories within 100 m around each lake (buffer zone) was calculated in QGIS 3.4.1 with GRASS 7.4.2 using ATKIS^®^ land use data with a 10 × 10 meter grid scale (© GeoBasis-DE/BKG 2013; AdV - Working Committee of the Surveying Authorities of the States of the Federal Republic of Germany, 2006). The ATKIS^®^-object categories were merged to seven land use classes: (1) urban (all anthropogenic infrastructures like buildings, streets, railroad tracks etc.), (2) agriculture (all arable land like fields and orchards but not meadows or pastures), (3) forest, (4) wetland (e.g., swamp lands, fen, peat lands), (5) excavation (e.g., open pit mine), (6) water (e.g., lakes, rivers, canals) and (7) other (not fitting in previous classes like succession areas, grass land, boulder sites etc.).

### Recreational use intensity

The lake-specific recreational use intensity was assessed by counting the type and number of recreationists during each site visit (between six and nine visits per lake, see biodiversity sampling below). Indirect use intensity metrics encompassed measures of accessibility and litter, which were assessed as follows: the pathways around every lake were enumerated with a measuring wheel (NESTLE-Cross-country Model 12015001, 2 m circumference, 0.1% accuracy), measuring the length of all trails and paths at each lake. These variables were summed and normalized to shoreline length. Angling sites and other open spaces accessible to other recreationists (e.g., swimmers) along the shoreline were counted, and all litter encountered along paths and sites was counted and assigned to (1) angling-related (e.g., lead weight, nylon line, artificial bait remains) and (2) other litter not directly angling-related (e.g., plastic packaging, beer bottles, cigarette butts). More intensively used lakes were expected to receive large amounts of litter and be more easily accessible through paths and trampled sites, which could affect biodiversity negatively.

### Age and morphology

The age of each lake was assessed through records in the angling clubs and by interviewing owners of lakes and regional administration or municipalities. Bathymetry and the size of each lake was mapped with a SIMRAD NSS7 evo2 echo sounder paired with a Lawrence TotalScan transducer mounted on a boat driving at 3 – 4 km/h along transects spaced at 25-45 m depending on lake size and depth. The data were processed using BioBase (Navico), and the post-processed data (depth and gps-position per ping) were used to calculate depth contour maps using ordinary kriging with the gstat-package in R version 3.5.1 (Gräler, Pebesma, & Heuvelink, 2016; R Core Team, 2013). Maximum depth and relative depth ratio (Damnjanović et al., 2018) were extracted from the contour maps. Shoreline length and lake area were estimated in QGIS 3.4.1 and used to calculate the shoreline development factor (*SDF*), which is the ratio of the lakes shoreline length (*L*) to the circumference of a circle with the same area (*A*):

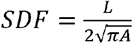

### Water chemistry and nutrient levels

During spring overturn, epilimnic water samples were taken to analyze total phosphorus concentrations (TP), total organic carbon (TOC), ammonium and nitrate concentrations (NH_4_, NO_3_) and chlorophyll a (CHLa; three samples per lake) as a measure of algal biomass. TP was determined using the ammonium molybdate spectrophotometric method (EN ISO 6878, 2004; Murphy & Riley, 1962), TOC was determined with a nondispersive infrared detector (NDIR) after combustion (DIN EN 1484, 1997), ammonium and nitrate were assessed using the spectrometric continuous flow analysis (DIN EN ISO 13395, 1996; EN ISO 11732, 2005), and CHLa was quantified using high performance liquid chromatography (HPLC), where the phaeopigments (degradation products) were separated from the intact chlorophyll a and only the concentration of the latter was measured (Mantoura & Llewellyn, 1983; Wright, 1991). For CHLa the mean of three samples per lake was determined for each sampling. Also during spring overturn, each lake’s conductivity and pH were measured in epilimnic water with a WTW Multi 350i sensor probe (WTW GmbH, Weilheim, Germany), and turbidity was assessed using a standard Secchi-disk. For all variables, the mean values of two years (i.e. two samplings) were used in the analyses.

### Littoral and riparian habitat assessment

Riparian structures and littoral dead wood were assessed using a plot design inspired by Kaufmann & Whittier (1997). Each plot consisted of a 15 × 4 m riparian sub-plot, a 1 × 4 m shoreline band, and a 4-meter-wide littoral transect extending into the lake to a maximum of ten meters or a water depth of three meters. At each lake the position of the first plot was randomly selected and subsequent plots placed every 100 (or 150 m for larger lakes) apart along the shoreline until the lake was surrounded, resulting in 4-20 plots per lake (depending on lake size). In each riparian sub-plot and shoreline band, all plant structures (e.g., trees, tall herbs, reed) were assessed following the protocol of Kaufmann & Whittier (1997) as 0 - absent, 1 - sparse (< 10 % coverage), 2 - moderate (10-39 % coverage), 3 - dominant (40-75 % coverage), and 4 - very dominant (> 75 % coverage). In each littoral transect all dead wood was counted, and length and bulk diameters measured. Additionally, the width and height of each coarse woody structure was assessed, and each piece assigned to either (1) simple dead wood (bulk diameter < 5 cm and length < 50 cm, no or very low complexity), or (2) coarse woody structure (bulk diameter > 5 cm and/or length > 50, any degree of complexity) following the criteria of DeBoom & Wahl (2013). Further, for each dead wood structure the volume was calculated using the formula for a cylinder for simple dead wood and for an ellipsoid for coarse woody structure.

### Riparian plant species

All lakes were sampled for riparian plant species at four transects (one per cardinal direction) in May. Each transect was 100 m long and contained five evenly spaced (20 m distance) 1 m^2^-plots. Along the transects trees (>2 m high) were identified following Spohn, Golte-Bechtle, & Spohn (2015) and counted. Within each sampling plot, riparian vascular plants (<2 m high) were identified following the same key (Spohn et al., 2015) and their abundance assessed following Braun-Blanquet (1964). The regional species pool was estimated from the Red Lists of Lower Saxony (Garve, 2004), which includes a full species inventory, in combination with their expected occurrence according to habitat type and species’ habitat preferences.

### Submerged macrophytes

All lakes were sampled for submerged macrophytes between late June and late August, following the sampling protocol of Schaumburg, Schranz, Stelzer, & Vogel (2014). Every lake was scuba dived and snorkeled along transects set perpendicular to the shoreline from the bank (depth = 0 m) to the middle of the lake until the deepest point of macrophyte growth was reached. The position of the first transect was randomly chosen and all other transects spaced evenly along the shoreline at 80-150 m distances depending on lake size, resulting in 4-20 transects sampled per lake. Along each transect, in every depth stratum (0-1 m, 1-2 m, 2-4 m, 4-6 m) the dominance of submerged macrophyte species was visually estimated following the Kohler scale (Kohler, 1978). No macrophytes were found below 6 m depth. Macrophytes were identified directly under water or if this was not possible, samples were taken and identified under a stereomicroscope following Van de Weyer & Schmitt (2011). Stonewort species were only identified to the genus level (*Chara* and *Nitella*), thus exact species numbers might be underestimated. Macrophyte dominance was transformed to percent coverage for each transect (Van der Maarel, 1979). The average coverage per stratum was extrapolated to the total lake using the contour maps. The total macrophyte coverage in the littoral zone was calculated using the extrapolated coverage from strata between 0 m and 3 m depth. The regional species pool was estimated from the Red Lists of Lower Saxony, which include full species inventories, in combination with the expected species for gravel pit lakes following the list of plant species associations in Lower Saxony (Garve, 2004; Korsch, Doege, Raabe, & van de Weyer, 2013; Preising et al., 1990).

### Amphibians

Amphibians were sampled during the mating-seasons (from March to May). Every lake was sampled twice: (1) during the day with an inflatable boat driving slowly along the shore searching for adults, egg-balls (frogs) and egg-lines (toads), (2) after sunset by foot around the lake searching for calling adults. Each observation (adult or eggs) was marked with a GPS (Garmin Oregon 600) and identified in the field or photographed for later identification following Schlüpmann (2005). Numbers were recorded (adults) or estimated (eggs), assuming 700 to 1500 eggs per egg-ball (frogs) or 10,000 eggs per (100 % covered) m^2^ of egg-line-assemblages (toads). The egg numbers were calculated from pictures taken in the field and verified with literature (Trochet et al., 2014). The regional species pool was estimated from the Red List of Lower Saxony, which includes a full species inventory, in combination with their expected distribution (Podloucky & Fischer, 2013).

### Odonata

Dragonflies and damselflies were sampled once per lake between early- and mid-summer. At each lake, the whole shoreline was intensively searched during mid-day. Sitting or flushing adult individuals were caught with a hand net (butterfly net, 0.2 mm mesh size, bioform), identified using Lehmann & Nüss (2015), and released without being harmed. The regional species pool was estimated from the Red List of Lower Saxony, which includes a full species inventory, in combination with their expected habitat preferences (Altmüller & Clausnitzer, 2010; Lehmann & Nüss, 2015).

### Waterfowl and songbirds

Waterfowl were identified following Dierschke (2016) and counted at every visit (between six and nine visits per lake). Songbirds were sampled once per lake between early- and mid-summer using a point-count sampling combined with a bioacoustics approach which was also used in other studies (Rempel, Hobson, Holborn, Van Wilgenburg, & Elliott, 2005; Wilson, Barr, & Zagorski, 2017). Two-minutes audio-recordings were taken (ZOOM Handy Recorder H2, Surround 4-Channel setting, 44.1kHz sampling frequency, 16 bit quantification) at sampling points placed 200 m apart around the whole lake, assuming each sampling point covers a radius of 100 m. Sampling points were marked with GPS. At each point all birds seen (or heard while not recording) were also noted when identified following Dierschke (2016). The audio-records were analyzed in the lab, and singing species were identified using reference audio samples (www.deutsche-vogelstimmen.de; www.vogelstimmen-wehr.de) and a birdsong-identifying software (BirdUp - Automatic Birdsong Recognition, developed by Jonathan Burn, Version 2018). The regional species pools for waterfowl and songbirds were estimated from the Red List of Lower Saxony (Krüger & Nipkow, 2015), which includes a full species inventory, in combination with their expected occurrence according to habitat type and preferences (Dierschke, 2016).

### Diversity metrics

The analysis focused on species presence-absence data to arrive at measures of taxonomic species richness, an aggregate index of species diversity. Additionally, the Simpson diversity index (Pielou, 1969) was computed using relative abundance data by species to consider the dominance of certain species within each taxa-specific community. There was no consideration in whether a particular species detected actually recruits in a given gravel pit lake, rather only consideration in the species inventory present, assuming the estimates represented a minimal estimate of local richness as rare species likely remained undetected. To weigh rare and threatened species more heavily, additionally the richness of threatened species was computed, and an index of taxon-specific conservation value for the study region was estimated following Oertli et al. (2002). To that end, each species was ranked according to its threat status on the Red Lists of Lower Saxony (Altmüller & Clausnitzer, 2010; Garve, 2004; Korsch et al., 2013; Krüger & Nipkow, 2015; Podloucky & Fischer, 2013). Species of least concern were ranked lowest (*c*(0) = 2° = 1). All species classified with an increasing threat status category r according to the regional Red List were weighted exponentially more strongly as *c*(*r*) = 2^*r*^ (Table 2) following Oertli et al. (2002). For each lake, the final taxon-specific conservation value (*CV*) was calculated as the sum of all values for the observed species *s*_*i*_ (*s*_1_, *s*_2_, *s*_3_,…, *s*_*n*_) divided by the total number of species (*n*) for a given taxon:

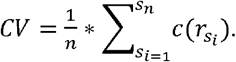

**Table 2:**
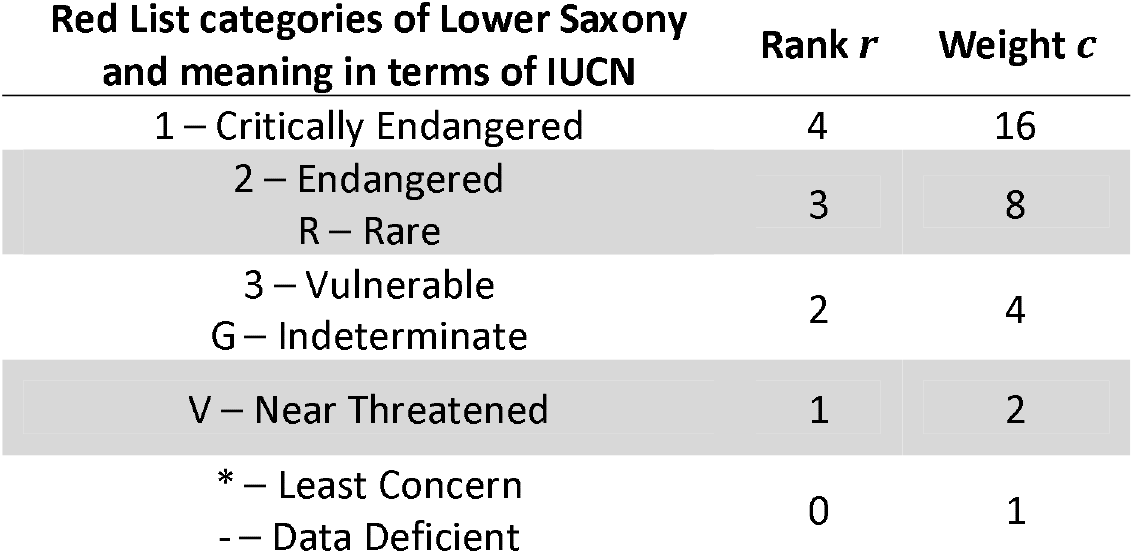
Ranking of Red List categories used for calculation of conservation values.

The conservation index value increases with more species of a given taxon being threatened or rare. A range of different allocations of threat status were tested to estimate the conservation value, using also national and European red lists; however, the results remained robust. For space reasons, just the regional index is reported here.

Finally, to test for differences in species composition across all lakes, the pooled species inventory by lake type (managed and unmanaged) was used, and the Sørensen index (Sørensen, 1948) as a measure of community similarity was calculated. The Sørensen index ranges from 0 (no species in common among the two lakes types) to 1 (all species the same) and is calculated as 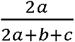, with *a* being the number of shared species and b and c being the numbers of unique species to each lake type, respectively. As an indicator for whether species compositions are substantial (i.e., biologically meaningful) different or not, so called “faunal brakes” as well as “floral breaks” were searched for. Following Matthews (1986) faunal or floral brakes among lake types were assumed to occur when the Sørensen index was < 0.5.

### Statistical analysis

The impact of the presence of recreational-fisheries management on aquatic and riparian biodiversity was tested in two steps.

First, differences in taxon-specific species richness, Simpson diversity index, richness of threatened species, conservation value and as well as key environmental variables between lake types (managed and unmanaged gravel pits) were assessed with univariate statistics. To that end, mean differences among lake types were tested using Student’s t (in case of variance homogeneity) or Welch-F (in case of variance heterogeneity) whenever the error term was normally distributed (Shapiro-Wilk-test). Otherwise, a Mann-Whitney-U-test of median differences was used. P-values were Sidak-corrected (Šidák, 1967) for multiple comparisons. Significance was assessed at *P* < 0.05.

Second, the among-lake variation in species richness was modelled as a function of lake type and a set of lake-specific environmental descriptors. These analyses aimed at further isolating an impact of fisheries management and type of recreational uses on species inventory across all taxa and lakes in a joint model that included other predictor variables of the lake environment. To reduce the dimensionality of the environmental variables, multiple Principal Component Analyses (PCA) without rotations were conducted on related classes of environmental variables respectively (groups were structured into variables related to morphology, productivity, habitat structure, land use, and recreational use). Environmental variables forming Principal Components (PC) were considered correlated, their loadings identified, the axis meanings interpreted, and the PC scores used as indicator variables. A multivariate Redundancy Analyses (RDA) was then conducted to examine if recreational fisheries management explains variation in either environmental variables or species richness across multiple taxa in the multivariate space. In addition to lake type, all relevant environmental variables (e.g., trophic state, surface area/steepness, land use, riparian/littoral habitat structure, water chemistry), recreational use intensity, gravel pit age, and catchment were included in the multivariate analysis of species richness. With the RDA, a forward selection process (Blanchet, Legendre, & Borcard, 2008) was used to identify the environmental predictors that explained the most variance in species richness across different taxa and lakes, including management as a key variable of interest in this study. Using the variance inflation factor (VIF; Neter, Kutner, Nachtsheim, & Wasserman, 1996) correlated environmental variables were removed before model building. All data were scaled and centered (z-transformation) prior to analyses. The degree of explanation was expressed using the adjusted coefficient of multiple determinations (*R*^2^_adj_). Variables significantly explaining variation in richness across lakes were also assessed using ANOVA (Analysis of variance) at a significance level of *P* < 0.05. All calculations and analyses were carried out in R version 3.5.1 using the vegan-package (Oksanen et al., 2018; R Core Team, 2013).

## 3. Results

### Description of lake types in relation to the environment

The studied lakes were, on average, small (mean ± SD, area 6.5 ± 5.2 ha, range 0.9 – 19.5 ha), shallow (maximum depth 9.6 ± 5.2 m, range 1.1 – 23.5 m) and mesotrohic (TP 26.3 ± 30.9 µg/l, range 8 - 160 µg/l) with moderate visibility (Secchi depth 2.4 ± 1.4 m, range 0.5 – 5.5 m) (Table 3). The land use in a 100 m buffer around the lake was, on average, characterized by low degree of forestation (mean 16 ± 21 %, range 0 – 72.6 %) and high degree of agricultural land use (mean 27 ± 22 %, range 2.4 – 79 %). Lakes were, on average, situated close to both human settlements (mean distance to the next village 618.3 ± 533.4 m, range 20 – 1810 m) and other water bodies (mean distance to next lake, river, or canal 55.8 ± 84.7 m, range 1 – 305 m). Gravel pit lakes were all in an advanced stage of succession and on average 27.3 ± 13.3 years old (range 6 – 54 years, see Tables S1-S4 for detailed lake-specific environmental variables). The study lakes belong to four different catchments (small North Sea tributaries and the catchments of the rivers Ems, Weser and Elbe; Table 1, Figure 1).

**Table 3:** Univariate comparison of environmental variables between managed and unmanaged gravel pit lakes. *P*-values are Sidak-corrected to account for multiple comparisons within classes of related environmental variables, **significant ones** (***P* < 0.05) are bolded**, statistical trends (*P* < 0.1) are in italics.

| Class | Environmental variable (abbreviation) | mean ± standard deviation<br>(minimum - maximum) | | Comparison | | |
| --- | --- | --- | --- | --- | --- | --- |
|  |  | Managed<br>(N = 16) | Unmanaged<br>(N = 10) | Test * | Statistic | *P*-value |
| Morphology | Maximum depth (m)<br>(MaxDep) | 9.7 ± 4.9<br>(2.8-23.5) | 9.5 ± 6.0<br>(1.1-23.0) | U-test | W = 89 | 0.986 |
| | Lake area (ha)<br>(LArea) | 7.4 ± 5.6<br>(1.0-19.5) | 4.9 ± 4.2<br>(0.9-13.6) | U-test | W = 105 | 0.592 |
| | Shoreline development factor<br>(SDF) | 1.5 ± 0.3<br>(1.1-2.2) | 1.6 ± 0.3<br>(1.3-2.2) | t-test | t = -1.0 | 0.824 |
| | Relative depth ratio<br>(RelDepR) | 0.04 ± 0.01<br>(0.02-0.07) | 0.04 ± 0.02<br>(0.01-0.07) | t-test | t = -1.1 | 0.754 |
| Trophic state | Total phosphorus ($\mu\text{g} \cdot \text{l}^{-1}$)<br>(TP) | 25.7 ± 36.5<br>(8-160) | 27.2 ± 20.9<br>(12-72) | U-test | W = 63.5 | 0.983 |
| | Total organic carbon ($\text{mg} \cdot \text{l}^{-1}$)<br>(TOC) | 6.6 ± 2.8<br>(2.5-13) | 6.0 ± 2.5<br>(2.9-12.4) | U-test | W = 93 | 0.997 |
| | Mean chlorophyll a ($\mu\text{g} \cdot \text{l}^{-1}$)<br>(CHLa) | 11.6 ± 15.9<br>(2.05-65.3) | 21.2 ± 26.1<br>(2.6-90.6) | U-test | W = 53 | 0.765 |
| | Secchi depth (m)<br>(Secchi) | 2.6 ± 1.5<br>(0.5-5.5) | 2.0 ± 1.1<br>(0.5-4.5) | t-test | t = 0.9 | 0.972 |
| | Ammonium ($\mu\text{g} \cdot \text{l}^{-1}$)<br>(NH4) | 56.9 ± 71.2<br>(15.0-240.0) | 84.0 ± 164.8<br>(15.0-550.0) | U-test | W = 75 | 1.000 |
| | Nitrate ($\mu\text{g} \cdot \text{l}^{-1}$)<br>(NO3) | 283.8 ± 380.4<br>(5.0-1040.0) | 733.0 ± 984.1<br>(5.0-2940.0) | U-test | W = 50 | 0.604 |
| | Conductivity ($\mu\text{S} \cdot \text{cm}^{-1}$)<br>(Con) | 450.9 ± 192.3<br>(136-742) | 478.6 ± 279.8<br>(159-957) | t-test | t = -0.3 | 1.000 |
| | pH Value<br>(pH) | 7.9 ± 0.5<br>(6.7-9) | 8.1 ± 0.5<br>(7.5-9.2) | t-test | t = -1 | 0.967 |
| Habitat structure | Volume-% of simple dead wood<br>(SDW_Vol) | 0.005 ± 0.009<br>(0-0.035) | 0.008 ± 0.010<br>(0.001-0.028) | U-test | W = 65.5 | 0.973 |
| | Volume-% of coarse woody structure<br>(CWS_Vol) | 1.5 ± 1.4<br>(0.02-5.6) | 1.7 ± 2.0<br>(0.02-6.2) | U-test | W = 86.5 | 1.000 |
| | Mean riparian tree cover on an ordinal<br>scale from 0 to 4 (Rip_Trees) | 1.0 ± 0.2<br>(0.4-1.5) | 0.9 ± 0.3<br>(0.4-1.2) | t-test | t = 0.9 | 0.942 |
| | Mean litoral reed cover on an ordinal<br>scale from 0 to 4 (Reed) | 1.3 ± 0.9<br>(0-2.5) | 0.8 ± 0.7<br>(0-1.7) | t-test | t = 1.25 | 0.779 |
| | Mean riparian vascular plants cover on an<br>ordinal scale from 0 to 4 (Rip_Herbs) | 1.7 ± 0.4<br>(1.1-3.0) | 1.0 ± 0.7<br>(0.1-1.9) | U-test | W = 118 | 0.225 |
|  | *Submerged macrophyte cover in the*<br>*littoral zone in % (MP_Cov)* | *39.3 ± 19.9*<br>*(12.5-82.3)* | *21.1 ± 27.5*<br>*(0-85.2)* | *U-test* | *W = 126* | *0.083* |
| Age | Lake age in years by 2017<br>(Age) | 29.4 ± 12.4<br>(11-52) | 23.8 ± 14.7<br>(6-54) | t-test | t = 1.05 | 0.303 |

\* t-test = Student's t-test / U-test = Mann-Whitney-U-test

### Environmental characteristics of managed and unmanaged gravel pit lakes

Both lake types did not statistically differ in age, size, trophic state, and land use (Table 3). A similar result was revealed in a multivariate RDA, which confirmed the absence of significant differences between managed and unmanaged lakes in “classes of environmental variables” (i.e., PC scores, for details see Tables S5, S6) representing morphology (an index of steepness and water body size; *R*^2^_adj_ = −0.005, F = 0.86, P = 0.470), trophic state (*R*^2^_adj_ = −0.006, F = 0.86, P = 0.544), proximity to alternative water bodies (*R*^2^_adj_ = −0.023, F = 0.45, P = 0.867), proximity to human presence (*R*^2^_adj_ = −0.035, F = 1.90, P = 0.143) and land use variables (*R*^2^_adj_ = −0.033, F = 1.85, P = 0.135). However, in multivariate space the habitat structure differed significantly among managed and unmanaged lakes along the first PC axis (Dim 1), which represented a vegetation gradient below and above water (Figure 2). Along this axis, managed lakes were found to be more vegetated than unmanaged ones in both the riparian and the littoral zones (*R*^2^_adj_ = −0.056, F = 2.48, P = 0.022).

**Figure 2:**
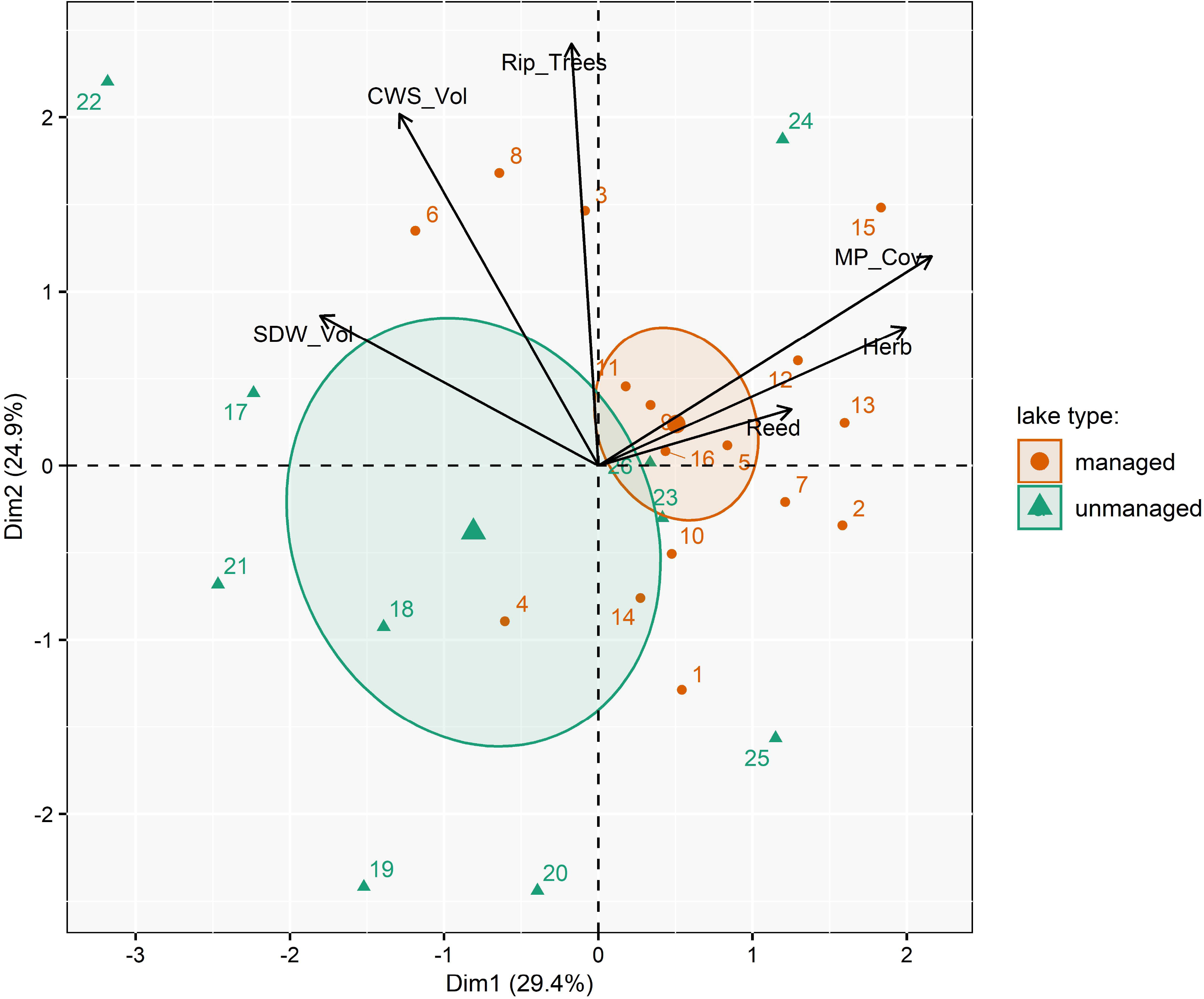
Principal component analysis (PCA) by classes of related environmental variables visualized for habitat structure (SDW_Vol = volume-% of simple dead wood, CWS_Vol = volume-% of coarse woody structure, Rip_Trees = mean riparian tree cover, Herb = mean riparian vascular plants cover, Reed = mean litoral reed cover, MP_Cov = submerged macrophyte cover in the littoral zone; Table 3). Percentages in brackets show the proportional variance explained by each axis respectively. Numbers reflect the different lakes (Table 1). The centroids of lake types are plotted as supplementary variables that did not influence the ordination. The 95% confidence-level around centroids are plotted to visualize differences between lake types.

### Recreational uses of managed and unmanaged lakes

The two lake types differed strongly in terms of recreational use intensity, particularly in relation to the observed angling intensity. As intended by the study design, managed lakes revealed, on average, significantly higher angling use intensity indexed by a diverse set of variables like angling litter density, extension of open sites, paths and trails and number of anglers observed (Table 4). By contrast, the average recreational use intensity of managed and unmanaged lakes by non-angling recreationists (e.g., swimmers) did not statistically differ when analyzed by univariate statistics on a variable-by-variable level (Table 4). However, when all indicator variables of the recreational use, both angling and non-angling, were combined in a multivariate RDA analysis as a function of lake type, managed lakes separated from unmanaged lakes along PC axis 1. This axis represented differences in recreational use intensity by both anglers and other recreationists (particularly swimmers) and extension of trails and paths (Figure 3, *R*^2^_adj_ = −0.16, F = 5.76, P < 0.001). Note that there was no differentiation among lake types along the second PC axis of the recreational variables (Figure 3), which represented shoreline (in)accessibility.

**Table 4:** Univariate comparison of environmental variables between managed and unmanaged gravel pit lakes. *P*-values are Sidak-corrected to account for multiple comparisons within classes of related environmental variables, **significant ones** (***P* < 0.05) are bolded**, statistical trends (*P* < 0.1) are in italics.

| Class | Environmental variable<br>(abbreviation) | mean ± standard deviation<br>(minimum - maximum) | | Comparison | | |
| --- | --- | --- | --- | --- | --- | --- |
|  |  | Managed<br>(N = 16) | Unmanaged<br>(N = 10) | Test * | Statistic | *P*-value |
| Land use | Excavation within 100m-buffer (%) (Excav) | 4.8 ± 7.2<br>(0-21.3) | 9.4 ± 14.6<br>(0-39.0) | U-test | W = 63 | 0.718 |
| | Agriculture within 100m-buffer (%) (Agric) | 22.8 ± 19.9<br>(2.4-55.9) | 33.8 ± 25.4<br>(3.5-79.0) | U-test | W = 58 | 0.598 |
| | Forest within 100m-buffer (%) (Forest) | 22.0 ± 25.1<br>(0-72.6) | 5.6 ± 6.0<br>(0-15.5) | U-test | W = 118 | 0.132 |
| | Urban area within 100m-buffer (%) (Urban) | 27.8 ± 29.2<br>(0-87.5) | 17.4 ± 24.5<br>(0-59.5) | U-test | W = 99 | 0.767 |
| Water | Wetland within 100m-buffer (%) (Wetland) | 0.9 ± 3.6<br>(0-14.4) | 5.1 ± 14.1<br>(0-45.1) | U-test | W = 61.5 | 0.504 |
| | Water surface within 100m-buffer (%) (Water) | 9.7 ± 12.5<br>(0.9-50.4) | 8.7 ± 8.8<br>(0.5-30.1) | U-test | W = 80 | 1.000 |
| | Distance to next lake (m) (DistLake) | 164.1 ± 236.4<br>(5-850) | 264.1 ± 440.6<br>(1-1280) | U-test | W = 87 | 0.999 |
| | Distance to next river (m) (DistRiver) | 5226.1 ± 9805.2<br>(25-29,900) | 3999.5 ± 9841.6<br>(220-31,920) | U-test | W = 92 | 0.980 |
| | Distance to next canal (m) (DistCanal) | 312.4 ± 462.3<br>(1-1630) | 224.5 ± 367.9<br>(5-1180) | U-test | W = 84 | 1.000 |
| Human presence | Distance to next road (m) (DistRoad) | 265.3 ± 314.4<br>(15-1010) | 558.0 ± 510.1<br>(30-1530) | U-test | W = 50.5 | 0.416 |
| | Distance to next settlement (m) (DistVille) | 504.1 ± 407.8<br>(20-1400) | 801.0 ± 673.0<br>(60-1810) | t-test | t = -1.4 | 0.530 |
| | Distance to next city (m) (DistCity) | 7135.0 ± 4087.6<br>(170-13,130) | 5859.0 ± 4488.3<br>(1070-15,110) | t-test | t = 0.8 | 0.917 |
| Recreational use | Litter related to angling (pieces per m shoreline) (A_Lit) | 0.05 ± 0.05<br>(0-0.20) | 0.002 ± 0.007<br>(0-0.021) | U-test | W = 140.5 | **0.007** |
| | Litter unrelated to angling (pieces per m shoreline) (NonA_Lit) | 0.70 ± 0.50<br>(0.02-1.48) | 0.34 ± 0.71<br>(0-2.29) | U-test | W = 126 | 0.124 |
| | Angling-sites and open spaces (% of shoreline) (open_sites) | 18.5 ± 19.8<br>(3.6-87.7) | 8.4 ± 14.4<br>(0-48.6) | U-test | W = 133 | **0.044** |
| | Trails and paths per shoreline (m*m<sup>-1</sup>) (Trails) | 0.9 ± 0.1<br>(0.6-1.1) | 0.4 ± 0.5<br>(0-1.4) | U-test | W = 138 | **0.019** |
| | Anglers per lake per visit (Anglers) | 1.6 ± 1.6<br>(0-5.1) | 0.1 ± 0.2<br>(0-0.8) | U-test | W = 143 | **0.006** |
| | Dog walkers per lake per visit (Dogs) | 1.7 ± 1.9<br>(0-6) | 0.5 ± 1.0<br>(0-3.3) | U-test | W = 123.5 | 0.154 |
| | Swimmers per lake per visit (Swimmers) | 2.9 ± 2.6<br>(0-10) | 0.7 ± 1.0<br>(0-3.1) | U-test | W = 129.5 | 0.075 |
| | Other recreationists per lake per visit (other_people) | 2.9 ± 3.2<br>(0.3-11.9) | 0.9 ± 1.4<br>(0-3.8) | U-test | W = 128.5 | 0.087 |

\* t-test = Student's t-test / U-test = Mann-Whitney-U-test

**Figure 3:**
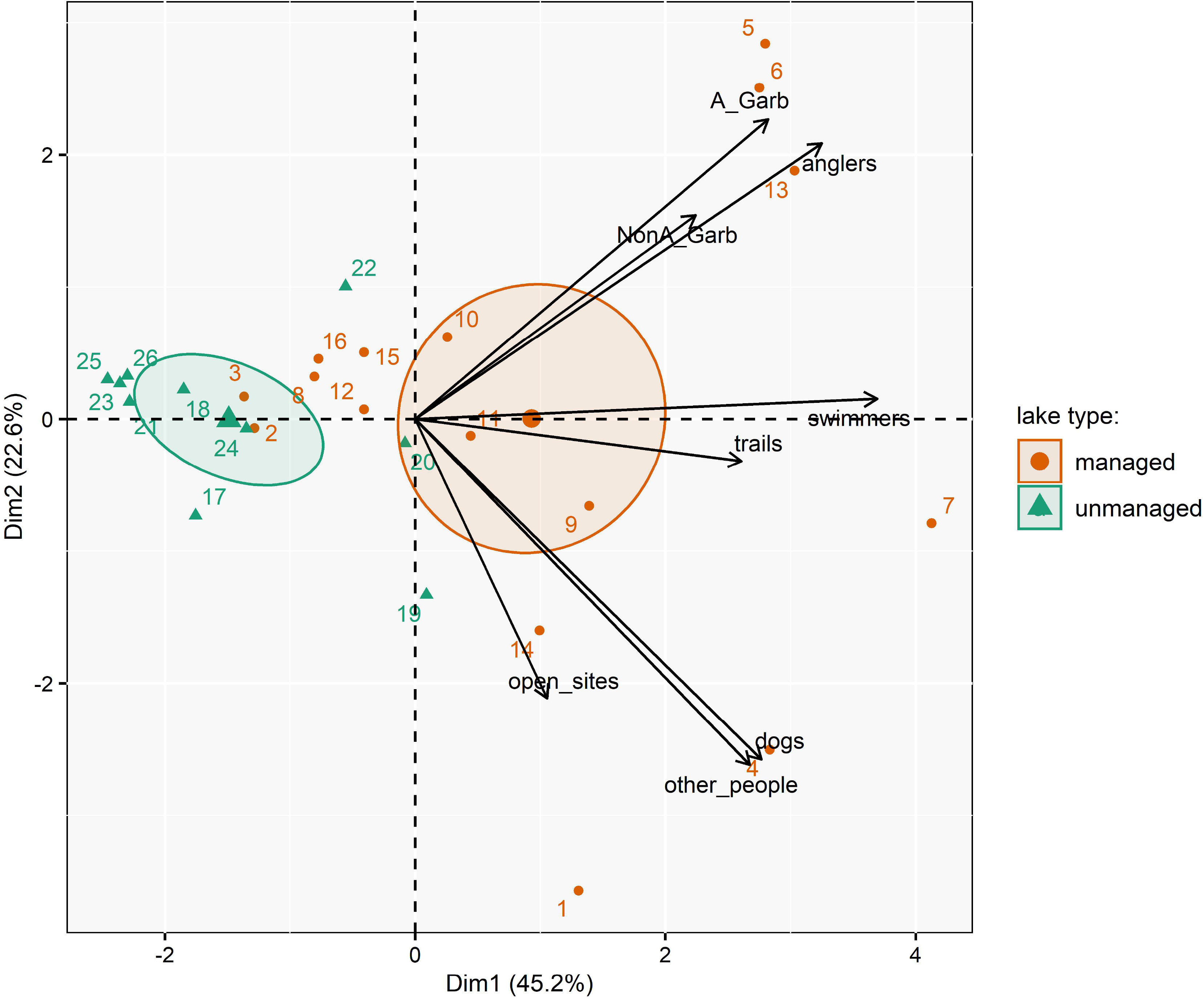
Principal component analysis (PCA) by classes of related environmental variables visualized for recreational use intensity (A_Lit = litter related to angling, NonA_Lit = litter unrelated to angling, open_sites = angling-sites and open spaces, Trails = trails and paths per shoreline, Anglers = no. per visit, Dogs = dog walkers per visit, Swimmers = no. per visit, other_people = other recreationists per visit; Table 4). Percentages in brackets show the proportional variance explained by each axis respectively. Numbers reflect the different lakes (Table 1). The centroids of lake types are plotted as supplementary variables that did not influence the ordination. The 95% confidence-level around centroids are plotted to visualize differences between lake types.

### Species diversity and taxon-specific conservation value in managed and unmanaged gravel pit lakes

In total 41 submerged macrophytes species were detected, 191 riparian vascular plants, 44 trees, 3 amphibians, 33 Odonata, 36 songbirds and 34 waterfowl species. This species inventory represented a substantial fraction of the regional species pool of trees (59 %), Odonata (56 %) and waterfowl (45 %). By contrast, only one third or less of the regional species pool of amphibians (38 %), songbirds (33 %), submerged macrophytes (33 %), and vascular plant species (12 %) were detected. Only few species non-native to Lower Saxony or Germany were found: four submerged macrophyte species (e.g., *Elodea nuttallii*, [Planch.] H. St. John, which is invasive), four riparian tree species, two waterfowl species (e.g., *Alopochen aegyptiaca*, L., which is invasive), one riparian vascular plant species, and one dragonfly species.

Based on the pooled species inventories (gamma diversity), unique species (i.e. species present in only one lake or only one lake type) were found in all taxonomic groups except amphibians (Table 5). Managed lakes hosted more unique species within most taxonomic groups than unmanaged lakes, while unmanaged lakes had more unique Odonata. No faunal or floral breaks were detected between managed and unmanaged lakes using the Sørensen index (all indices ≥ 0.5; Table 5). The average taxon-specific species richness (alpha-diversity), the Simpson diversity index, the average number of threatened species and the average taxon-specific conservation value were statistically similar in managed and unmanaged lakes across all taxonomic groups when analyzed using univariate statistics (Table 6).

**Table 5:**
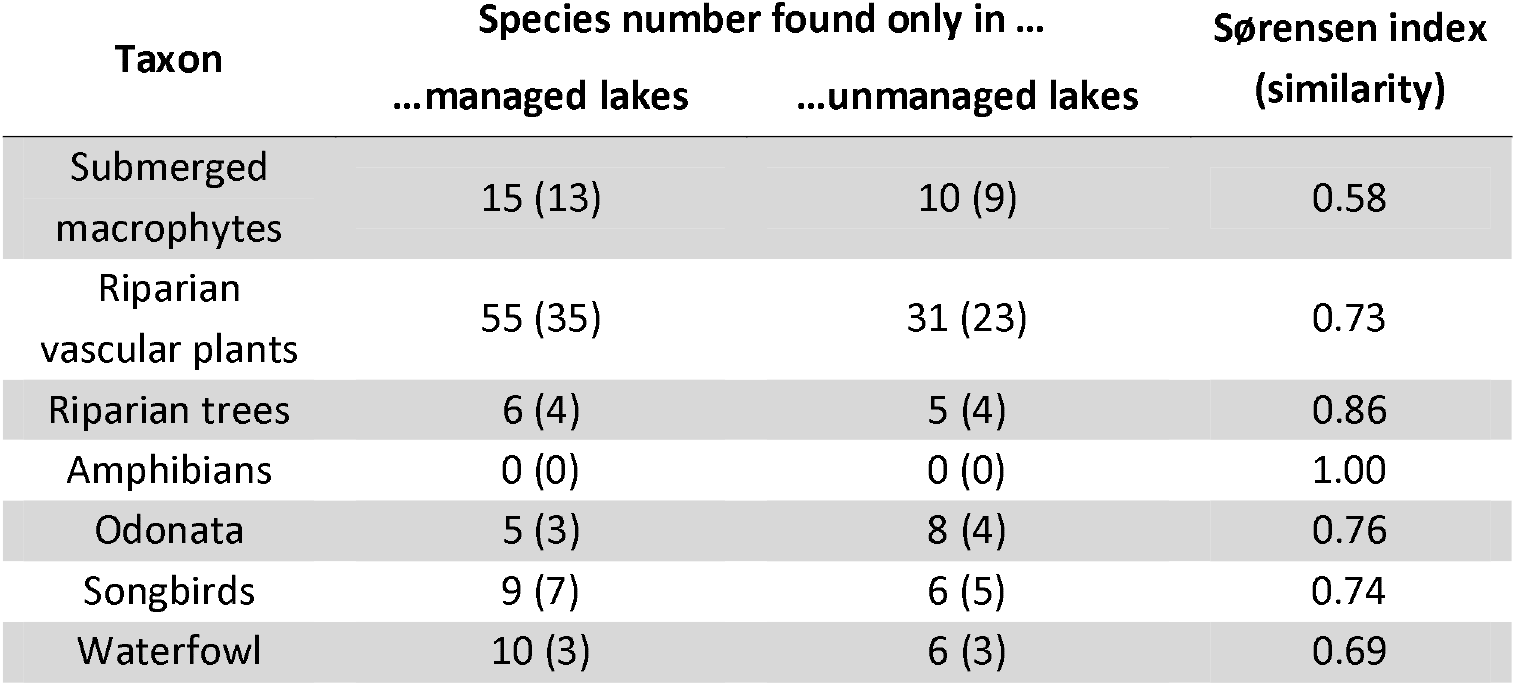
Overview about unique species of different taxa found at managed and unmanaged gravel pits in Lower Saxony, Germany. The numbers in brackets refer to single-lake observations, i.e. the number of species found at only one lake each.

**Table 6:**
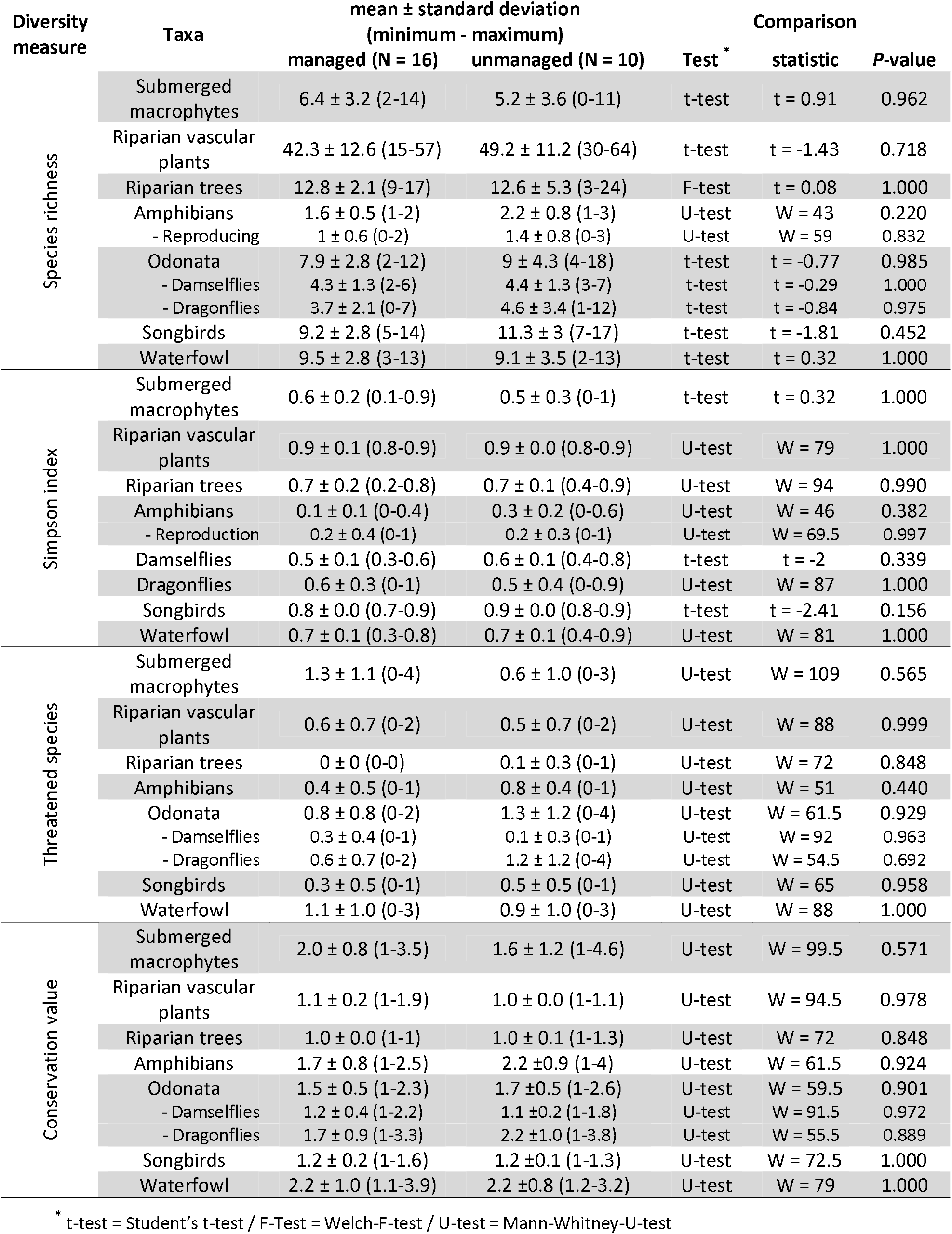
Univariate comparison of species richness, Simpson index, threatened species and taxon-specific conservation values in managed and unmanaged gravel pit lakes. **Statistical differences of Sidak-corrected *P*-values < 0.05 are bolded**.

### Environmental correlates of among-lake variation in species richness

Across lakes, species richness of amphibians, Odonata, songbirds and riparian vascular plant species covaried along the first axis (Figure 4), collectively representing riparian diversity (for full PCA results, see Table S8). The second PCA axis represented mainly submerged macrophytes (Figure 4). The third axis was related to the diversity of riparian tree species and the forth mainly to waterfowl diversity (Figure 5). Therefore, lakes offering high riparian species richness were not necessarily rich in biodiversity of submerged macrophytes, waterfowl or trees. The RDA analysis to explain the among-lake variation in species richness as a function of lake type alone revealed no influence of this factor on among-lake richness across several taxa (RDA, *R*^2^_adj_ = 0.028, F = 1.73, P = 0.114).

**Figure 4:**
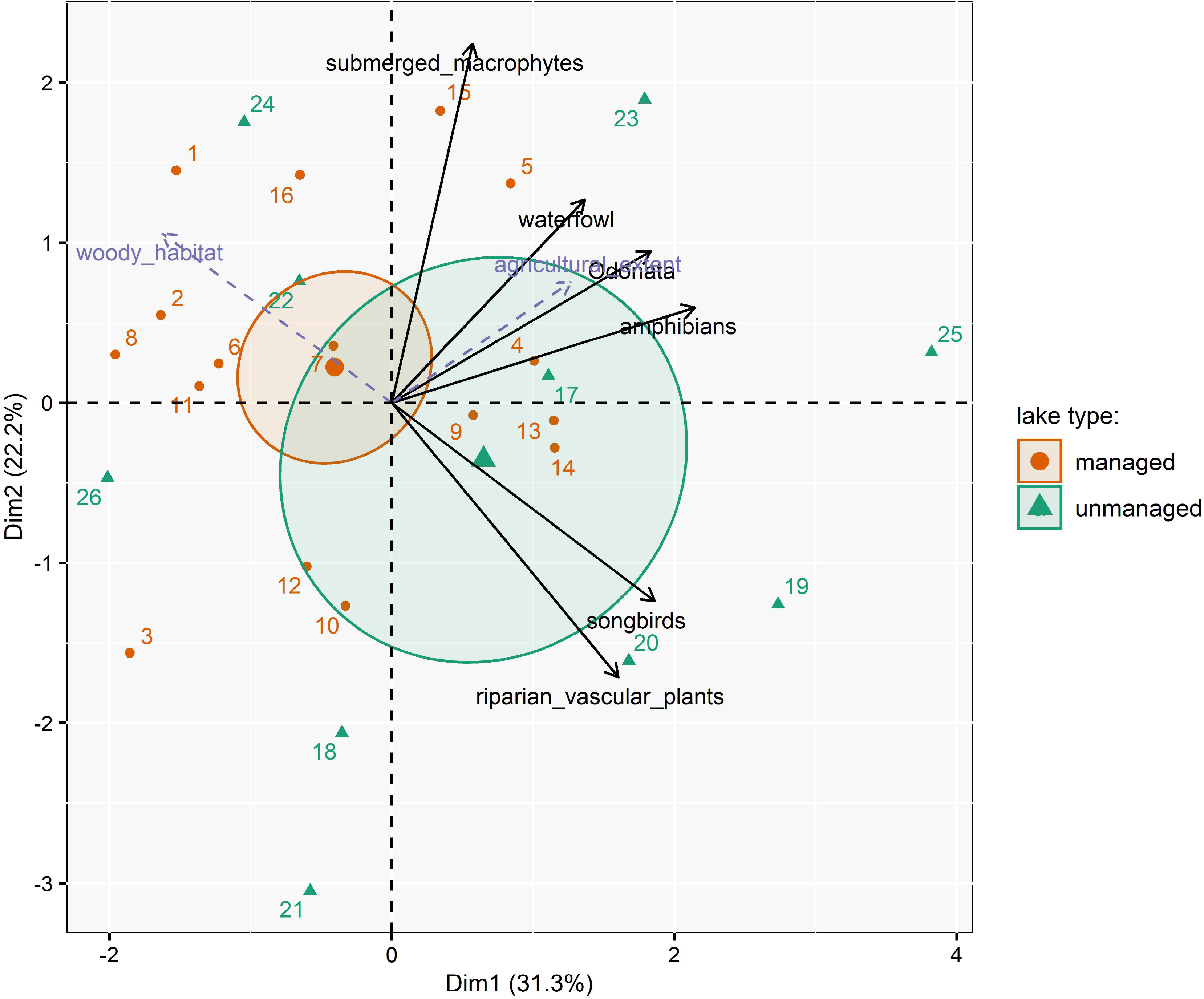
Principal component analysis (PCA) of species richness plotted for the first two axes (only relevant, i.e. highly contributing, variables are shown). Percentages in brackets show the proportional variance explained by each axis respectively. Numbers reflect the different lakes (Table 1). The centroids of lake types and the explanatory variables from redundancy analysis (RDA, slashed purple lines, only the important ones for Dim1 and Dim2 are shown) are plotted as supplementary variables to not influence the ordination. The 95% confidence-level around centroids are plotted to visualize differences between lake types.

**Figure 5:**
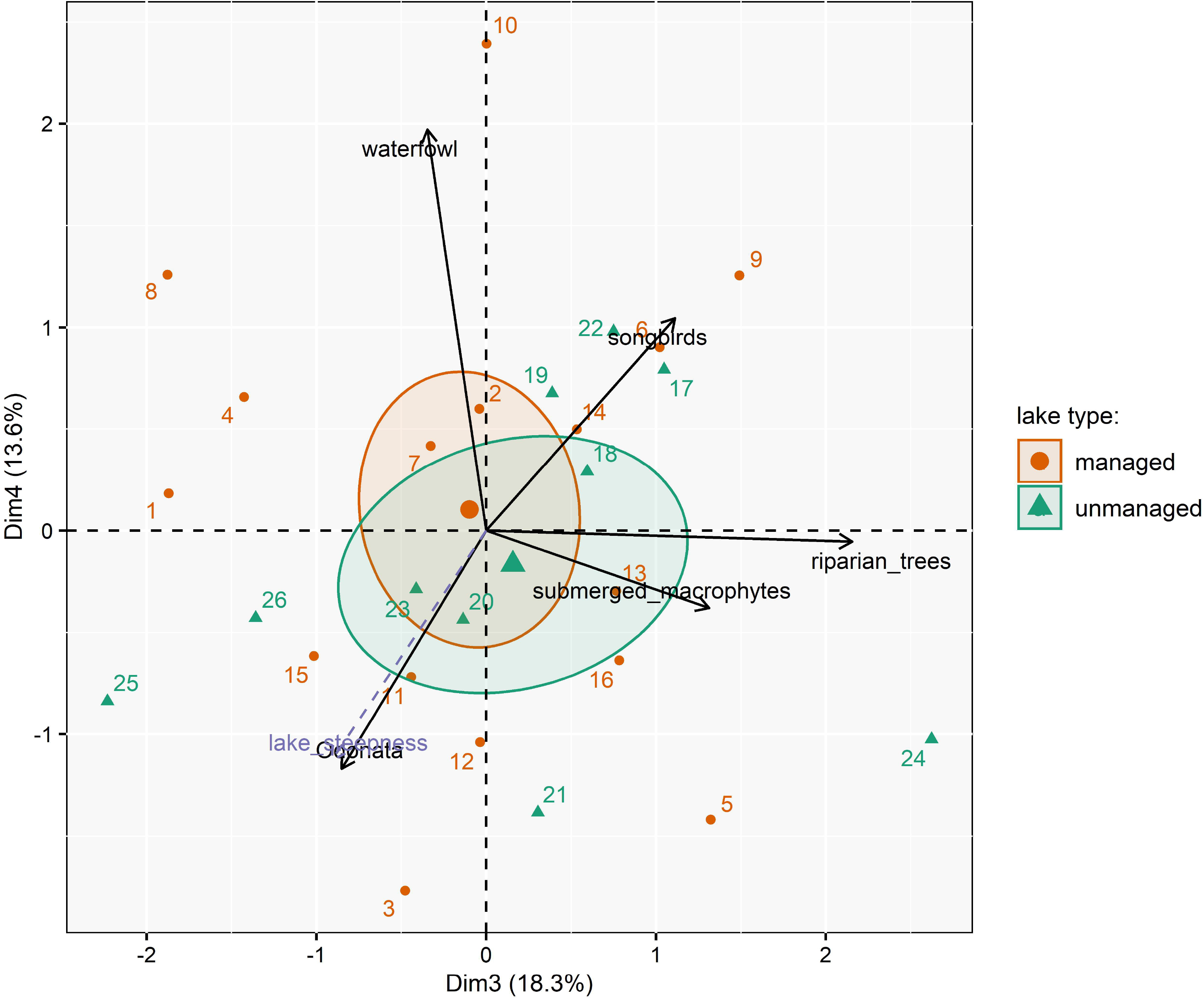
Principal component analysis (PCA) of species richness plotted for the third and fourth axis (only relevant, i.e. highly contributing, variables are shown). Percentages in brackets show the proportional variance explained by each axis respectively. Numbers reflect the different lakes (Table 1). The centroids of lake types and the explanatory variables from redundancy analysis (RDA, slashed purple lines, only the important ones for Dim3 and Dim4 are shown) are plotted as supplementary variables to not influence the ordination. The 95% confidence-level around centroids are plotted to visualize differences between lake types.

All environmental variables subsumed by PC-scores into environmental predictors and lake age had acceptable inflation factors (VIF < 5, maximum: 4.98, supplement Table S7) and were used along with catchment association and lake type in the full RDA analysis to explain among-lake species richness jointly across all taxa. The RDA-based forward model selection retained a few environmental variables as key correlates of species richness of multiple taxa across lakes, but lake type dropped from the best model (Table 7). Therefore, among lake variation in richness across several aquatic and riparian taxa was solely explained by environmental factors unrelated to either lake type or recreation-related variables. Specifically, the coverage of woody habitat along the littoral was negatively correlated with riparian species richness and positively correlated with tree diversity along the first axis in Figure 4. The extent of agricultural land use (representing also more rural conditions, supplementary Table S6) was positively associated with riparian species richness (Figure 4). Lake steepness (representing also small lake size and low shoreline development factor, supplementary Table S5) was negatively correlated with waterfowl species richness (Figure 5). All other environmental variables, including lake age and catchment were not significant (Table 7). The best model explained 36 % of the total variance in the multivariate species richness. In this model, neither lake type nor any of the recreational use variables explained variation in species richness of a range of aquatic and riparian taxa among lakes.

**Table 7:**
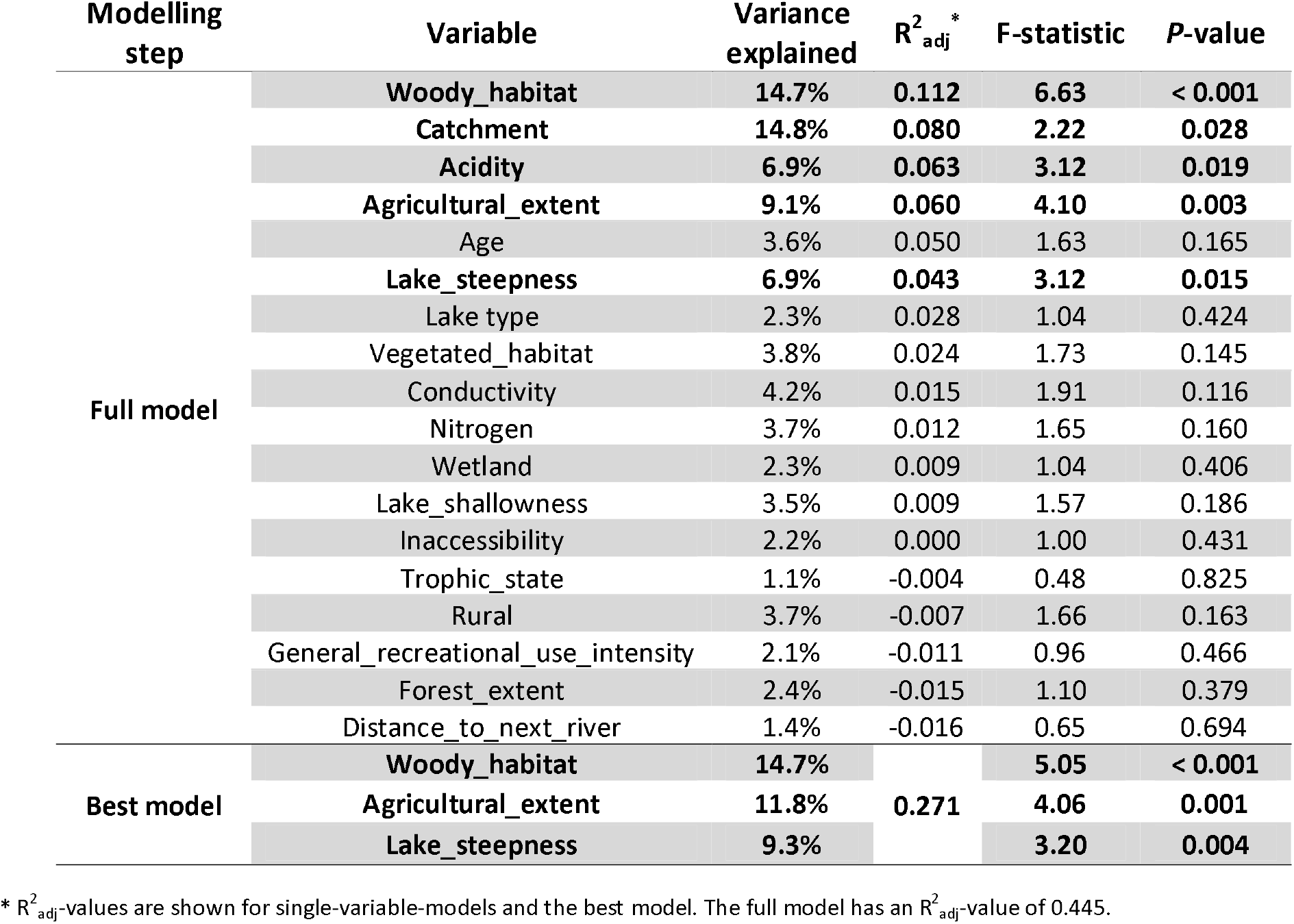
ANOVA results of forward selection of RDA models explaining species richness across taxa. Variables are ordered by their *R*^2^_adj_-value. **Significant variables (*P* < 0.05) are bolded**.

| Modelling step | Variable | Variance explained | $R^2_{adj}$ * | F-statistic | P-value |
| --- | --- | --- | --- | --- | --- |
| Full model | **Woody_habitat** | **14.7%** | **0.112** | **6.63** | **< 0.001** |
|  | **Catchment** | **14.8%** | **0.080** | **2.22** | **0.028** |
|  | **Acidity** | **6.9%** | **0.063** | **3.12** | **0.019** |
|  | **Agricultural_extent** | **9.1%** | **0.060** | **4.10** | **0.003** |
|  | Age | 3.6% | 0.050 | 1.63 | 0.165 |
|  | **Lake_steepness** | **6.9%** | **0.043** | **3.12** | **0.015** |
|  | Lake type | 2.3% | 0.028 | 1.04 | 0.424 |
|  | Vegetated_habitat | 3.8% | 0.024 | 1.73 | 0.145 |
|  | Conductivity | 4.2% | 0.015 | 1.91 | 0.116 |
|  | Nitrogen | 3.7% | 0.012 | 1.65 | 0.160 |
|  | Wetland | 2.3% | 0.009 | 1.04 | 0.406 |
|  | Lake_shallowness | 3.5% | 0.009 | 1.57 | 0.186 |
|  | Inaccessibility | 2.2% | 0.000 | 1.00 | 0.431 |
|  | Trophic_state | 1.1% | -0.004 | 0.48 | 0.825 |
|  | Rural | 3.7% | -0.007 | 1.66 | 0.163 |
|  | General_recreational_use_intensity | 2.1% | -0.011 | 0.96 | 0.466 |
|  | Forest_extent | 2.4% | -0.015 | 1.10 | 0.379 |
|  | Distance_to_next_river | 1.4% | -0.016 | 0.65 | 0.694 |
| Best model | **Woody_habitat** | **14.7%** |  | **5.05** | **< 0.001** |
|  | **Agricultural_extent** | **11.8%** | **0.271** | **4.06** | **0.001** |
|  | **Lake_steepness** | **9.3%** |  | **3.20** | **0.004** |

\* $R^2_{adj}$ -values are shown for single-variable-models and the best model. The full model has an $R^2_{adj}$ -value of 0.445.

## 4. Discussion

In line with initial expectations no differences in species richness, Simpson diversity and conservation value were found across all examined taxa among managed and unmanaged gravel pit lakes, and a similar species pool was found to be present in both lake types. Collectively, this study did not reveal that recreational-fisheries management (through impacts on fish communities) or the presence of anglers (through disturbance effects on shoreline habitat and wildlife or lethal impacts through lost fishing gear) significantly constrains the development of diverse communities of amphibians, birds, submerged macrophytes, terrestrial plants and Odonata relative to those expected at lakes that are not managed for recreational fisheries. Instead, the best predictors of the variation in species richness among lakes were found to be related to land use variables, the extent of woody habitat at the lake shores and the lake morphology (size and steepness). Therefore, this study suggests that for the taxonomic groups and lake types that were examined broader environmental factors and land use, and not the presence of recreational fisheries and its management of fish stocks and littoral zones, shape taxonomic diversity of plants, birds, amphibians, and dragonflies.

### Biodiversity potential of gravel pit lakes

Gravel pit lakes in Lower Saxony, Germany, were found to host a substantial species diversity and fraction of the regional species’ pools of aquatic and riparian taxa, in particular trees, Odonata and waterfowl. This finding supports related work in other areas of Europe (Damnjanović et al., 2018; Oertli, 2018; Spyra & Strzelec, 2019). Yet, only small fractions of the regional species’ pools were detected for vascular plant species, submerged macrophytes, songbirds and amphibians. In particular amphibians are considered very sensitive to predation from fish (Hecnar & M’Closkey, 1997) and none of our study lakes were free of fish (Matern et al., 2019). Many amphibian species depend on shallow water and best develop in small, temporary waters (Shulse, Semlitsch, Trauth, & Williams, 2010). The gravel pits of this study were also relatively steeply-sloped with small fractions of littoral areas, disconnected from rivers, placed in agricultural landscapes and close to anthropogenic infrastructure. All of these factors are negative for amphibian diversity and can explain the low species richness detected for this taxon (Shulse et al., 2010). Importantly, the results of this study indicate that management by recreational fisheries and the substantially different fish communities in managed and unmanaged lakes can be excluded as an additional stressor.

### Environmental differences among managed and unmanaged lakes

The gravel pit lakes studied were similar in the majority of the environmental factors examined (including age) except the coverage of submerged macrophytes, which was more prevalent in managed gravel pit lakes compared to unmanaged ones. Submerged macrophytes have been reported to be strongly affected by stocking of benthivorous fishes, such as common carp (Bajer et al., 2016; Miller & Crowl, 2006). However, in a subset of the same gravel pit lakes presented here, Matern et al. (2019) found similar biomasses of fishes in managed and unmanaged lakes, with carps and breams being present in both lake types. Due to the sampling gear used by Matern et al. (2019), the authors likely underestimated the abundance and biomass of common carp and other large benthivorous fish (Ravn et al., 2019). Although no absolute biomass data of carp or other species in the study lakes are available, the fact that submerged macrophytes were more diverse and more developed in the angler-managed lakes suggests that co-existence of carp and other game fish with a species rich submerged macrophyte community, also in terms of threatened stonewort species (*Chara sp*., *Nitella sp*.), is possible. This disagrees with expectations expressed elsewhere that managing lakes with benthivorous fish necessarily harms submerged macrophytes (Van de Weyer, Meis, & Krautkrämer, 2015). Instead, the more developed submerged macrophytes in managed lakes revealed here suggests that critical biomass thresholds for benthivorous fish after which macrophytes often vanish or strongly decline (about 100 kg/ha; Vilizzi, Tarkan, & Copp, 2015) might not have been reached in the study lakes. Alternatively, the transferability of typical mesocosm studies that have reported substantial impacts of carp on macrophytes to occur after reaching about 100 kg/ha may not hold under conditions in the wild (Arlinghaus, Hühn, et al., 2017).

Lake shorelines managed by anglers were previously reported to be heavily modified to accommodate angling sites and access to anglers (O’Toole et al., 2009). At the same time, crowding is a severe constraint that reduces angler satisfaction (Beardmore, Hunt, Haider, Dorow, & Arlinghaus, 2015). Although improved accessibility in angler-managed lakes was supported in this study, the amount of aquatic and riparian vegetation was significantly larger in angler-managed systems compared to unmanaged lakes. This indicates that maintaining accessibility of lakeshores to anglers does not necessarily mean degraded riparian or littoral habitat quality. Anglers have an interest to maintain access to lakes to be able to fish, but there is also an interest in developing suitable habitats for fish (Meyerhoff et al., 2019) and maintaining sites that promise solitude during the experience (Beardmore et al., 2015), which indirectly may also support biodiversity. We speculate the regular shoreline development activities by anglers and angling clubs to maintain access to angling sites may create “disturbances” (O’Toole et al., 2009), that regularly interrupt the succession of tree stands, thereby reducing the shading effects in the littoral zone (Monk & Gabrielson, 1985), thereby promoting growth of submerged macrophytes (Holtmann, Kerler, Wolfgart, Schmidt, & Fartmann, 2019). The littoral zone is the most productive habitat of lakes (Winfield, 2004), and most fish species depend on submerged macrophytes and other structures for spawning, foraging and refuge (Lewin, Mehner, Ritterbusch, & Brämick, 2014). Therefore, although anglers regularly engage in shoreline development activities and angling site maintenance, the data of this study suggest they do so to a degree that may maintain or even foster aquatic and riparian vegetation.

### Differences in recreational use of managed and unmanaged lakes

Managed lakes were found to have more developed tracks, paths, parking places and other facilities that attract anglers and other recreationists. Thus, angler-managed lakes were generally more accessible to water-based recreationists, although these differences were not always statistically significant among the two lake types for recreational uses other than angling. Importantly, despite managed lakes receiving regular fisheries-management activities such as stocking and angler use, neither “lake type” nor the index of general recreational use intensity were related with species richness across multiple taxa and lakes. Thus, for the diversity metrics and the taxa that were examined (vegetation, Odonata, amphibians, birds), this study does not suggest that the use of gravel pits by recreational fisheries significantly constraints the development of aquatic and riparian biodiversity across a range of taxa. Clearly, species-specific effects on disturbance-sensitive species (e.g., selected bird species; Knight, Anderson, & Marr, 1991) may still occur, which the aggregate metrics of taxonomic richness or the Simpson community diversity index might have been too insensitive to detect. Further work on community differences among managed and unmanaged lakes is warranted.

### Differences in biodiversity among managed and unmanaged lakes

Across all taxa that were examined, no statistical differences were found in species richness, number of threatened species, conservation value and Simpson diversity between managed and unmanaged lakes. This result was unexpected. Recreational-fisheries management can affect aquatic and riparian biodiversity through various pathways, first through supporting and enhancing fish stocks that exert predation pressure (e.g., on tadpoles and Odonata larvae; Hecnar & M’Closkey, 1997; Knorp & Dorn, 2016), second through indirect fish-based effects (e.g., uprooting macrophytes through benthivorous feeding; Bajer et al., 2016), third through direct removal or damage of submerged and terrestrial plants during angling activities (O’Toole et al., 2009), which may have knock-on effects on dragonflies (Z. Müller et al., 2003), and fourth through activity-based disturbance effects or lethal impacts through lost fishing sinkers in particular on birds (Cryer et al., 1987; Sears, 1988). This study design was not tailored towards directly measuring disturbance effects on particular species; instead it was designed to examine a range of taxonomic richness indices in aggregate for communities present at gravel pit lakes that are subjected to recreational fisheries compared to ecologically similar lakes that do not. When judged against these aggregate biodiversity metrics, the study presented here does not support the idea that recreational-fisheries management and angler presence has important impacts that modify species inventories to a degree that strongly depart from situations expected at unmanaged lakes without anglers. Previous work has reported relevant reductions in bird biodiversity from lakes exposed to human disturbances due to recreation including angling (Bell et al., 1997). However, similar species richness and conservation value of both waterfowl and riparian songbirds in managed and unmanaged lakes were found in the present work. This does not exclude the possibility that for example the breeding success of specific disturbance-sensitive taxa might have been impaired in angler-managed lakes (Park, Park, Sung, & Park, 2006; Reichholf, 1988). But if such effects were present, they were not strong enough to substantially alter species richness, not to be confused with species identity. Overall, against the chosen metrics, the findings supported the initial hypothesis of no impacts from recreational fishing on non-targeted taxa in gravel pits situated in agricultural landscapes.

### Environmental determinants of aquatic and riparian biodiversity in gravel pit lakes

The species richness of different taxa did not uniformly vary among lakes, in contrast to a study of managed shallow ponds by Lemmens et al. (2013). While examining strictly aquatic taxa (zooplankton, submerged and emerged aquatic macrophytes, benthic invertebrates), Lemmens et al. (2013) revealed uniform responses in species richness across taxa and ponds in their study. The much broader trophic and habitat requirements of aquatic and riparian taxa examined here resulted in significantly more variable biotic responses. For example, lakes rich in riparian biodiversity were not necessarily rich in submerged macrophytes and waterfowl biodiversity. The reason was that the aquatic and riparian biodiversity responded to many variables beyond those measured within the lake. The multivariate analyses revealed that variation in species richness across multiple taxa was driven by structural variables such as habitat quality, lake morphometry (surface area and steepness) and land use in a buffer zone around the lake, but not by recreational use intensity or the presence of recreational-fisheries management activities. Thus, environmental factors unrelated to recreational fishing seem to overwhelm any specific impacts of angling, at least for the taxonomic diversity metrics and the taxa examined here.

Mosaics of different habitats (reeds, overhanging trees etc.) along the shoreline support species richness and diversity for most taxa (Kaufmann, Hughes, Whittier, Bryce, & Paulsen, 2014), and the presence of endangered biodiversity increase the recreational value of gravel pit lakes as perceived by anglers (Meyerhoff et al., 2019). Extended woody habitat both in water and particularly in the riparian zone was correlated with increased tree diversity, but reduced riparian species richness of vascular plants, amphibians, Odonata and songbirds. This might be explained by the shading effect of trees on non-woody vegetation (Monk & Gabrielson, 1985). Odonata, songbirds and amphibian species benefited from more vegetated littoral habitats, in agreement with previous work (Paracuellos, 2006; Remsburg & Turner, 2009; Shulse et al., 2010). The species richness of waterfowl was strongly governed by lake surface area and steepness of the littoral, with larger and shallower lakes having a higher waterfowl species richness, confirming earlier findings by Paszkowski & Tonn (2000). The three dominant waterfowl species (occurring on 85 % or more of sampled lakes) were either omnivorous (mallard, *Anas platyrhynchos*, L.) or herbivorous-invertivorous (common coot, *Fulica atra*, L., and tufted duck, *Aythya fuligula*, L.). In addition, 77 % of the lakes were used by grey goose (*Anser anser*, L.), which feeds on terrestrial plants. Thus, it can be concluded that the dominant waterfowl detected at the studied lakes benefit from submerged macrophytes or riparian plants, both found to be more abundant at managed lakes.

Collectively, the presented data does not support substantial negative impacts of recreational fisheries management on the species richness and community diversity of waterfowl and songbirds present at gravel pit lakes. In a related study from Welsh reservoirs Cryer et al. (1987) observed only distributional changes of waterfowl to the presence of anglers and no changes in abundance. Similarly negligible effects of anglers on piscivorous birds at Canadian natural lakes were reported by Somers, Heisler, Doucette, Kjoss, & Brigham (2015). Specifically for gravel pit lakes, Bell et al. (1997) failed to find evidence for impacts of recreational fishing on the community structure of waterfowl, although in particular diving waterfowl reduced their abundance during presence of anglers and other recreationists. In that study, similar to the presented one, habitat quality and lake size were more important for waterfowl diversity than the bank use by anglers, and their shoreline management rather supported grazing waterfowl by opening up sites (Bell et al., 1997). This does not mean that recreational fishing will not impact bird populations at all as for example the breeding success of certain disturbance sensitive species might still be impaired (Park et al., 2006; Reichholf, 1988). However, this study was not designed to examine the breeding success of particular species and rather focused on aggregate diversity metrics. Against these, this study did not reveal any significant disturbance effects caused by recreational fisheries.

### Limitations

The strength of the study design is the focus on multiple taxa, which is rare in the recreational ecology literature related to freshwaters. The limitations are that it was not focused on specific species, and the sampling design does not answer whether the detected mobile species (e.g., birds or Odonata) recruited in the study lakes or only used them temporarily as feeding or resting habitat. Moreover, because of adjustments in taxa-specific sampling schemes, seasonal taxa (amphibians, Odonata) might have been underestimated in the sampling and rare species might likely been missed (Yoccoz, Nichols, & Boulinier, 2001). However, even if this is the case the conclusions presented are robust because this systematic error affected both lake types compared here.

This study used a comparative approach where lakes were not randomly allocated to either angler-managed lakes or controls. All lakes sampled were from the same geographical area and the age of the lakes and the wider environmental factors were similar; thus, the key differences among lake types related to the presence of recreational fishing. Therefore, the design would have been able to differentiate strong angling-induced biodiversity effects would they exist in reality.

A further limitation is that the design did not include entirely unused lakes where recreation whatsoever is prohibited. The present data have to be interpreted against the possibility that gravel pits situated in reserves with strictly no human access might show higher species diversity than revealed in the sample lakes. All the lakes were situated in agricultural environments and all were exposed to a certain recreational use. Background disturbance (Liley, Underhill-Day, Panter, Marsh, & Roberts, 2015) might have affected the observed species pool, affecting the detectability of species in the study region. The conclusions of the present work are also confined to the environmental gradients that could be observed. For example, higher angler use intensities than found in the present work might reveal different results.

Finally, the recreational use intensity was mainly recorded during weekdays when the field visits were done. Thus, potential high intensity phases at weekends might be unrepresented. However, this would even strengthen our conclusions, if the real recreational use of managed lakes was well beyond what was considered in this analysis.

## Conclusions

The present study shows that recreational-fisheries management does not constrain the establishment of a rich biodiversity of aquatic and riparian taxa in and around gravel pit lakes relative to conditions expected at ecologically similar unmanaged lakes. Environmental variables related to habitat quality, land use and lake morphometry rather than recreation-related drivers were key in driving the species richness for multiple taxa across lakes. This study thus shows that co-existence of recreational fisheries and aquatic and riparian biodiversity of high conservation value and richness is possible, at least under the specific ecological conditions of gravel pit lakes in agricultural landscapes. From a conservation perspective, it is suggested that recreational fishing clubs increasingly rely on habitat enhancement activities to support both fish and other taxa present at gravel pit lakes. Moreover, development of diverse shorelines as well as the creation of more gently sloped littoral areas is recommended as actions to be completed during the creation of gravel pit lakes. If these actions are taken, bans on recreational fishing, are unlikely to produce further conservation benefits if the aim is to create high species diversity, independent of a specific species identity.

## Acknowledgements

We want to thank Sven Matern for his important help in every stage of this study and we thank Charlotte Robinchon for her productive dead-wood-sampling-internship. We also thank Leander Höhne, Jasper Münnich, Nicola Wegener, Jara Niebuhr, Natalie Arnold, Rachel Fricke and Stéphane Mutel for their help with field sampling and data analysis. We thank Christopher Monk for immense help on statistical analysis, R-coding and checking the language of the manuscript. We thank Pieter Lemmens, Miguel Palmer, Jörg Freyhof and Petr Zajicek for their advice during data processing, analysis and interpretation. We thank Sabine Hilt and Klaus van de Weyer for training and advices on macrophyte sampling and for help in the identification. We thank Tobias Goldhammer and the chemical lab of IGB for help with water samples. We specifically thank the work of Barbara Stein. We thank Jan Hallermann, Asja Vogt, Alexander Türck and Jürgen Schreiber for helping with material and equipment. Moreover, we thank Angelsportverein Leer u. Umgebung e.V., Bezirksfischereiverband für Ostfriesland e.V., Angler-Verein Nienburg e.V., ASV Neustadt am Rübenberge e.V., Fischereiverein Hannover e.V., Niedersächsisch-Westfälische Anglervereinigung e.V., Ralf Gerken, Heike Vullmer and the Stiftung Naturschutz im Landkreis Rotenburg (Wümme), Henning Scherfeld, FV Peine-Ilsede u. Umgebung e.V., SFV Helmstedt u. Umgebung e.V., Verein der Sportfischer Verden (Aller) e.V., Verein für Fischerei und Gewässerschutz Schönewörde u. Umgebung e.V., Steffen Göckemeyer, the Xella Kalksandsteinwerke Niedersachsen GmbH & Co. KG, Thomas Reimer, Melanie and Heinz H. Nordmeyer, Achaz von Hardenberg, Johann Augustin, Dieter Klensang, Elke Dammann, Cordula Stein and Holcim Germany, Matthias Emmrich and the Angler Association of Lower Saxony for participating in this study. The presented study was jointly financed by the German Federal Ministry of Education and Research (BMBF) and the German Federal Agency for Nature Conservation (BfN) with funds granted by the German Federal Ministry for the Environment, Nature Conservation and Nuclear Safety (BMU) within the BAGGERSEE-Projekt (grant number: 01LC1320A; www.baggersee-forschung.de). Additional funding came through the STÖRBAGGER-project via the Landesverband Sächsischer Angler e.V., the Landesfischereiverband Bayern e.V., and the Angler Association of Lower Saxony (www.ifishman.de/en/projects/stoerbagger/), the Stiftung Fischerei, Umwelt-und Naturschutz Deutschland (FUND Stiftung) and the BMBF through the Aquatag project (the German Federal Ministry of Education and Research (grant 01LC1826E). The funders had no role in the design, analysis and interpretation of the data. Finally, we thank the editor Philip J Boon, Lee Jackson and anonymous reviewers for constructive feedback on the manuscripts that substantially improved our presentation.

## References

AdV - Working Committee of the Surveying Authorities of the States of the Federal Republic of Germany. (2006). Section 5.4 - Explanations on ATKIS^®^. In S. Afflerbach & W. Kunze (Eds.), Documentation on the Modelling of Geoinformation of Official Surveying and Mapping (GeoInfoDok) (5.1, pp. 1–74). München, Germany.

Altmüller, R., & Clausnitzer, H.-J. (2010). Rote Liste der Libellen Niedersachsens und Bremens: 2. Fassung, Stand 2007. Informationsdienst Naturschutz Niedersachsen, 30, 211–238.

Arlinghaus, R. (2005). A conceptual framework to identify and understand conflicts in recreational fisheries systems, with implications for sustainable management. Aquatic Resources, Culture and Development, 1, 145–174.

Arlinghaus, R., Hühn, D., Pagel, T., Beck, M., Rapp, T., & Wolter, C. (2017). Fischereilicher Nutzen und gewässerökologische Auswirkungen des Besatzes mit Karpfen (Cyprinus carpio) in stehenden Gewässern: Ergebnisse und Schlussfolgerungen aktueller Ganzseeexperimente und Meta-Analysen. Fischerei Und Fischmarkt in Mecklenburg-Vorpommern, 1, 36–46.

Arlinghaus, R., Müller, R., Rapp, T., & Wolter, C. (2017). Nachhaltiges Management von Angelgewässern: Ein Praxisleitfaden (Band 30). Berlin, Germany: Leibniz-Institute of Freshwater Ecology and Inland Fisheries (IGB).

Bajer, P. G., Beck, M. W., Cross, T. K., Koch, J. D., Bartodziej, W. M., & Sorensen, P. W. (2016). Biological invasion by a benthivorous fish reduced the cover and species richness of aquatic plants in most lakes of a large North American ecoregion. Global Change Biology, 22, 3937–3947. https://doi.org/10.1111/gcb.13377

Beardmore, B., Hunt, L. M., Haider, W., Dorow, M., & Arlinghaus, R. (2015). Effectively managing angler satisfaction in recreational fisheries requires understanding the fish species and the anglers. Canadian Journal of Fisheries and Aquatic Sciences, 72, 500–513. https://doi.org/10.1139/cjfas-2014-0177

Bell, M. C., Delany, S. N., Millett, M. C., & Pollitt, M. S. (1997). Wintering waterfowl community structure and the characteristics of gravel pit lakes. Wildlife Biology, 3, 65–78. https://doi.org/10.2981/wlb.1997.009

Blanchet, F. G., Legendre, P., & Borcard, D. (2008). wory variables. Ecology, 89, 2623–2632.

Braun-Blanquet, J. (1964). Pflanzensoziologie: Grundzüge der Vegetationskunde. In Pflanr⍰ensor⍰iologie (3rd ed.). Wien, Austria: Springer. https://doi.org/10.1007/978-3-7091-4078-9

Chambers, J. M., & Hastie, T. J. (1992). Statistical models in S (Vol. 251). Pacific Grove, CA: Wadsworth & Brooks/Cole; Advanced Books & Software.

Council of the European Communities. (1992). Council Directive 92/43/EEC of 21 May 1992 on the conservation of natural habitats and of wild fauna and flora. Official Journal of the European Communities, L206, 7–50.

Cryer, M., Linley, N. W., Ward, R. M., Stratford, J. O., & Randerson, P. F. (1987). Disturbance of overwintering wildfowl by anglers at two reservoir sites in south wales. Bird Study, 34, 191–199. https://doi.org/10.1080/00063658709476961

Cucherousset, J., Boulêtreau, S., Azémar, F., Compin, A., Guillaume, M., & Santoul, F. (2012). “Freshwater killer whales”: Beaching behavior of an alien fish to hunt land birds. PLoS ONE, 7, 1–6. https://doi.org/10.1371/journal.pone.0050840

Damnjanović, B., Novković, M., Vesić, A., Živković, M., Radulović, S., Vukov, D., … Cvijanović, D. (2018). Biodiversity-friendly designs for gravel pit lakes along the Drina River floodplain (the Middle Danube Basin, Serbia). Wetlands Ecology and Management, 27, 1–22. https://doi.org/10.1007/s11273-018-9641-8

Dear, E. J., Guay, P.-J., Robinson, R. W., & Weston, M. A. (2015). Distance from shore positively influences alert distance in three wetland bird species. Wetlands Ecology and Management, 23, 315–318. https://doi.org/10.1007/s11273-014-9376-0

DeBoom, C. S., & Wahl, D. H. (2013). Effects of coarse woody habitat complexity on predator-prey interactions of four freshwater fish species. Transactions of the American Fisheries Society, 142, 1602–1614. https://doi.org/10.1080/00028487.2013.820219

Díaz, S., Settele, J., Brondizio, E. S., Ngo, H. T., Agard, J., Arneth, A., … Zayas, C. N. (2019). Pervasive human-driven decline of life on Earth points to the need for transformative change. Science, 1327. https://doi.org/10.1126/science.aaw3100

Dierschke, V. (2016). Welcher Vogel ist das? - 170 Vögel einfach bestimmen (3. Auflage). Stuttgart, Germany: Franckh-Kosmos.

DIN EN 1484. (1997). Water analysis - Guidelines for the determination of total organic carbon (TOC) and dissolved organic carbon (DOC).

DIN EN ISO 13395. (1996). Water quality - Determination of nitrite nitrogen and nitrate nitrogen and the sum of both by flow analysis (CFA and FIA) and spectrometric detection (ISO 13395:1996).

Dudgeon, D., Arthington, A. H., Gessner, M. O., Kawabata, Z.-I., Knowler, D. J., Lévêque, C., … Sullivan, C. A. (2006). Freshwater biodiversity: Importance, threats, status and conservation challenges. Biological Reviews of the Cambridge Philosophical Society, 81, 163–182. https://doi.org/10.1017/S1464793105006950

EN ISO 11732. (2005). Water quality - Determination of ammonium nitrogen - Method by flow analysis (CFA and FIA) and spectrometric detection (ISO 11732:1997).

EN ISO 6878. (2004). Water quality - Determination of phosphorus - Ammonium molybdate spectrometric method (ISO 6878:1998).

European Aggregates Association (UEPG). (2017). Estimates of aggregates production data 2017. Retrieved June 17, 2020, from http://www.uepg.eu/statistics/estimates-of-production-data/data-2017

Franson, J. C., Hansen, S. P., Creekmore, T. E., Brand, C. J., Evers, D. C., Duerr, A. E., & DeStefano, S. (2003). Lead fishing weights and other fishing tackle in selected waterbirds. Waterbirds: The International Journal of Waterbird Biology, 26, 345–352.

Garve, E. (2004). Rote Liste und Florenliste der Farn- und Blütenpflanzen in Niedersachsen und Bremen, 5. Fassung, Stand 1.3.2004. Informationsdienst Naturschutz Niedersachsen, 24, 1–76.

Gräler, B., Pebesma, E., & Heuvelink, G. (2016). Spatio-temporal interpolation using gstat. The R Journal, 8, 204–218.

Granek, E. F., Madin, E. M. P., Brown, M. A., Figueira, W., Cameron, D. S., Hogan, Z., … Arlinghaus, R. (2008). Engaging recreational fishers in management and conservation: Global case studies. Conservation Biology, 22, 1125–1134. https://doi.org/10.1111/j.1523-1739.2008.00977.x

Hecnar, S. J., & M’Closkey, R. T. (1997). The effects of predatory fish on amphibian species richness and distribution. Biological Conservation, 79, 123–131. https://doi.org/10.1016/s0006-3207(96)00113-9

Holtmann, L., Kerler, K., Wolfgart, L., Schmidt, C., & Fartmann, T. (2019). Habitat heterogeneity determines plant species richness in urban stormwater ponds. Ecological Engineering, 138, 434–443. https://doi.org/10.1016/j.ecoleng.2019.07.035

Kaufmann, P. R., Hughes, R. M., Whittier, T. R., Bryce, S. A., & Paulsen, S. G. (2014). Relevance of lake physical habitat indices to fish and riparian birds. Lake and Reservoir Management, 30, 177–191. https://doi.org/10.1080/10402381.2013.877544

Kaufmann, P. R., & Whittier, T. R. (1997). Habitat assessment. In J. R. Baker, D. V. Peck, & D. W. Sutton (Eds.), Environmental monitoring and assessment program – Surface waters: Field operations manual for lakes (pp. 5–1 to 5–26). Washington, D.C.: U.S. Environmental Protection Agency.

Knight, R. L., Anderson, D. P., & Marr, N. V. (1991). Responses of an avian scavenging guild to anglers. Biological Conservation, 56, 195–205. https://doi.org/10.1016/0006-3207(91)90017-4

Knorp, N. E., & Dorn, N. J. (2016). Mosquitofish predation and aquatic vegetation determine emergence patterns of dragonfly assemblages. Freshwater Science, 35, 114–125. https://doi.org/10.1086/684678

Kohler, A. (1978). Methoden der Kartierung von Flora und Vegetation von Süßwasserbiotopen. Landschaft + Stadt, 10, 73–85.

Korsch, H., Doege, A., Raabe, U., & van de Weyer, K. (2013). Rote Liste der Armleuchteralgen (Charophyceae) Deutschlands 3. Fassung, Stand: Dezember 2012. Hausknechtia, 17, 1–32.

Krüger, T., & Nipkow, M. (2015). Rote Liste der in Niedersachsen und Bremen gefährdeten Brutvogelarten, 8. Fassung, Stand 2015. Informationsdienst Naturschutz Niedersachsen, 35, 181–256.

Legendre, P., & Legendre, L. (2012). Numerical ecology (3rd Editio). Amsterdam, Netherlands: Elsevier.

Lehmann, A. W., & Nüss, J. H. (2015). Libellen - Bestimmungsschlüssel für Nord- und Mitteleuropa (6. Auflage). Göttingen, Germany: Deutscher Jugendbund für Naturbeobachtungen.

Lemmens, P., Mergeay, J., De Bie, T., Van Wichelen, J., De Meester, L., & Declerck, S. A. J. (2013). How to maximally support local and regional biodiversity in applied conservation? Insights from pond management. PLoS One, 8, e72538. https://doi.org/10.1371/journal.pone.0072538

Lewin, W.-C., Mehner, T., Ritterbusch, D., & Brämick, U. (2014). The influence of anthropogenic shoreline changes on the littoral abundance of fish species in German lowland lakes varying in depth as determined by boosted regression trees. Hydrobiologia, 724, 293–306. https://doi.org/10.1007/s10750-013-1746-8

Liley, D., Underhill-Day, J., Panter, C., Marsh, P., & Roberts, J. (2015). Morecambe Bay bird disturbance and access management report. Unpublished report by Footprint Ecology for the Morecambe Bay Partnership.

Mantoura, R. F. C., & Llewellyn, C. A. (1983). The rapid determination of algal chlorophyll and carotenoid pigments and their breakdown products in natural waters by reverse-phase high-performance liquid chromatography. Analytica Chimica Acta, 151, 297–314. https://doi.org/10.1016/S0003-2670(00)80092-6

Mardia, K. V., Kent, J. T., & Bibby, J. M. (1979). Multivariate Analysis. London, United Kingdom: Academic Press.

Matern, S., Emmrich, M., Klefoth, T., Wolter, C., Nikolaus, R., Wegener, N., & Arlinghaus, R. (2019). Effect of recreational-fisheries management on fish biodiversity in gravel pit lakes, with contrasts to unmanaged lakes. Journal of Fish Biology, 94, 865–881. https://doi.org/10.1111/jfb.13989

Matthews, W. J. (1986). Fish faunal “breaks” and stream order in the eastern and central United States. Environmental Biology of Fishes, 17, 81–92. https://doi.org/10.1007/BF00001739

McFadden, T. N., Herrera, A. G., & Navedo, J. G. (2017). Waterbird responses to regular passage of a birdwatching tour boat: Implications for wetland management. Journal for Nature Conservation, 40, 42–48. https://doi.org/10.1016/J.JNC.2017.09.004

Meyerhoff, J., Klefoth, T., & Arlinghaus, R. (2019). The value artificial lake ecosystems provide to recreational anglers: Implications for management of biodiversity and outdoor recreation. Journal of Environmental Management, 252, 109580. https://doi.org/10.1016/j.jenvman.2019.109580

Miller, S. A., & Crowl, T. A. (2006). Effects of common carp (Cyprinus carpio) on macrophytes and invertebrate communities in a shallow lake. Freshwater Biology, 51, 85–94. https://doi.org/10.1007/s00198-005-1915-3

Monk, C. D., & Gabrielson, F. C. J. (1985). Effects of shade, litter and root competition on old-field vegetation in South Carolina. Bulletin of the Torrey Botanical Club, 112, 383–392.

Müller, H. (2012). Zulässigkeit und Grenzen der Ausgestaltung/Einschränkung von Fischereirechten an Baggerseen [Rechtsgutachten]. Bayreuth/München, Germany: Bezirksfischereiverband Oberfranken e.V., Landesfischereiverband Bayern e.V.

Müller, Z., Jakab, T., Tóth, A., Dévai, G., Szállassy, N., Kiss, B., & Horváth, R. (2003). Effect of sports fisherman activities on dragonfly assemblages on a Hungarian river floodplain. Biodiversity and Conservation, 12, 167–179.

Murphy, J., & Riley, J. P. (1962). A modified single solution method for the determination of phosphate in natural water. Analytica Chimica Acta, 27, 31–36.

Neter, J., Kutner, M. H., Nachtsheim, C. J., & Wasserman, W. (1996). Applied linear statistical models (Fourth Edition). Chicago, IL: Irwin.

Nikolaus, R., Matern, S., Schafft, M., Klefoth, T., Maday, A., Wolter, C., … Arlinghaus, R. (n.d.). Impact of recreational fisheries management on the biodiversity of gravel pit lakes: a comparative study on multiple groups of aquatic organisms. Lauterbornia, in review.

O’Toole, A. C., Hanson, K. C., & Cooke, S. J. (2009). The effect of shoreline recreational angling activities on aquatic and riparian habitat within an urban environment: Implications for conservation and management. Environmental Management, 44, 324–334. https://doi.org/10.1007/s00267-009-9299-3

Oertli, B. (2018). Editorial: Freshwater biodiversity conservation: The role of artificial ponds in the 21st century. Aquatic Conservation: Marine and Freshwater Ecosystems, 28, 264–269. https://doi.org/10.1002/aqc.2902

Oertli, B., Joye, D. A., Castella, E., Juge, R., Cambin, D., & Lachavanne, J.-B. (2002). Does size matter? The relationship between pond area and biodiversity. Biological Conservation, 104, 59–70.

Oksanen, J., Blanchet, F. G., Friendly, M., Kindt, R., Legendre, P., McGlinn, D., … Wagner, H. (2018). vegan: Community Ecology Package (R package version 2.5-3).

Osgood, R. A. (2005). Shoreline density. Lake and Reservoir Management, 21, 125–126. https://doi.org/10.1080/07438140509354420

Paracuellos, M. (2006). Relationships of songbird occupation with habitat configuration and bird abundance in patchy reed beds. Ardea, 94, 87–98.

Park, J.-H., Park, H.-W., Sung, H.-C., & Park, S.-R. (2006). Effect of fishing activity on nest selection and density of waterfowls in Namyang Lake. Journal of Ecology and Field Biology, 29, 213–217.

Paszkowski, C. A., & Tonn, W. M. (2000). Effects of lake size, environment, and fish assemblage on species richness of aquatic birds. Internationale Vereinigung Für Theoretische Und Angewandte Limnologie: Verhandlungen, 27, 178–182. https://doi.org/10.1080/03680770.1998.11901222

Pielou, E. C. (1969). An introduction to mathematical ecology. New York City, NY: Wiley-Interscience.

Podloucky, R., & Fischer, C. (2013). Rote Listen und Gesamtartenlisten der Amphibien und Reptilien in Niedersachsen und Bremen, 4. Fassung. Informationsdienst Naturschutz Niedersachsen, 4.

Preising, E., Vahle, H.-C., Brandes, D., Hofmeister, H., Tüxen, J., & Weber, H. E. (1990). Die Pflanzengesellschaften Niedersachsens - Bestandsentwicklung, Gefährdung und Schutzprobleme: Salzpflanzengesellschaften der Meeresküste und des Binnenlandes // Wasserund Sumpfpflanzengesellschaften des Süßwassers. In B. Pilgrim (Ed.), Naturschutz Landschaftspflege Niedersachsens (20/7 & 8). Hannover, Germany: Niedersächsisches Landesamt für Ökologie - Naturschutz -.

R Core Team. (2013). R - A language and environment for statistical computing (3.5.1). Wien, Austria: R Foundation for Statistical Computing.

Randler, C. (2006). Disturbances by dog barking increase vigilance in coots Fulica atra. European Journal of Wildlife Research, 52, 265–270. https://doi.org/10.1007/s10344-006-0049-z

Ravn, H. D., Lauridsen, T. L., Jepsen, N., Jeppesen, E., Hansen, P. G., Hansen, J. G., & Berg, S. (2019). A comparative study of three different methods for assessing fish communities in a small eutrophic lake. Ecology of Freshwater Fish, 28, 341–352. https://doi.org/10.1111/eff.12457

Reichholf, J. H. (1988). Effects of anglers on the breeding of water birds in the internationally important wetland “Lower Inn River.” Vogelwelt, 109, 206–221.

Reid, A. J., Carlson, A. K., Creed, I. F., Eliason, E. J., Gell, P. A., Johnson, P. T. J., … Cooke, S. J. (2019). Emerging threats and persistent conservation challenges for freshwater biodiversity. Biological Reviews, 94, 849–873. https://doi.org/10.1111/brv.12480

Rempel, R. S., Hobson, K. A., Holborn, G., Van Wilgenburg, S. L., & Elliott, J. (2005). Bioacoustic monitoring of forest songbirds: interpreter variability and effects of configuration and digital processing methods in the laboratory. Journal of Field Ornithology, 76, 1–11. https://doi.org/10.1648/0273-8570-76.1.1

Remsburg, A. J., & Turner, M. G. (2009). Aquatic and terrestrial drivers of dragonfly (Odonata) assemblages within and among north-temperate lakes. Journal of the North American Benthological Society, 28, 44–56. https://doi.org/10.1899/08-004.1

Riedmüller, U., Hoehn, E., & Mischke, U. (2013). Auswerte-Tool für die Trophie-Klassifikation von Seen. Trophie-Index nach LAWA. (Computer Software; Version 1.0). Freiburg/Berlin, Germany: Limnologie-Büro Hoehn/Leibniz-Institute of Freshwater Ecology and Inland Fisheries (IGB).

Saulnier-Talbot, É., & Lavoie, I. (2018). Uncharted waters: The rise of human-made aquatic environments in the age of the “Anthropocene.” Anthropocene, 23, 29–42. https://doi.org/10.1016/j.ancene.2018.07.003

Schaumburg, J., Schranz, C., Stelzer, D., & Vogel, A. (2014). Verfahrensanleitung für die ökologische Bewertung von Seen zur Umsetzung der EG-Wasserrahmenrichtlinie: Makrophyten & Phytobenthos (Version 10). Augsburg/Wielenbach, Germany: Länderarbeitsgemeinschaft Wasser (LAWA).

Schlüpmann, M. (2005). Bestimmungshilfen: Faden- und Teichmolch-Weibchen, Braunfrösche, Wasser- oder Grünfrösche, Eidechsen, Schlingnatter und Kreuzotter, Ringelnatter- Unterarten. Rundbrief zur Herpetofauna von Nordrhein-Westfalen, 28, 1–38.

Sears, J. (1988). Regional and seasonal variations in lead poisoning in the Mute Swan (Cygnus olor) in relation to the distribution of lead and lead weights, in the Thames area, England. Biological Conservation, 46, 115–134.

Shulse, C. D., Semlitsch, R. D., Trauth, K. M., & Williams, A. D. (2010). Influences of design and landscape placement parameters on amphibian abundance in constructed wetlands. Wetlands, 30, 915–928. https://doi.org/10.1007/s13157-010-0069-z

Šidák, Z. (1967). Rectangular confidence regions for the means of multivariate normal distributions. Journal of the American Statistical Association, 62, 626–633. https://doi.org/10.1080/01621459.1967.10482935

Somers, C. M., Heisler, L. M., Doucette, J. L., Kjoss, V. A., & Brigham, R. M. (2015). Lake use by three avian piscivores and humanslll : Implications for angler perception and conservation. Journal of Open Ornithology, 8, 10–21.

Sørensen, T. A. (1948). A method of establishing groups of equal amplitude in plant sociology based on similarity of species content and its application to analyses of the vegetation on Danish commons. Biologiske Skrifter, 5, 1–34.

Spohn, M., Golte-Bechtle, M., & Spohn, R. (2015). Was blüht denn da? (59. Auflag). Stuttgart, Germany: Franckh-Kosmos.

Spyra, A., & Strzelec, M. (2019). The implications of the impact of the recreational use of forest mining ponds on benthic invertebrates with special emphasis on gastropods. Biologia, 1–12. https://doi.org/10.2478/s11756-019-00221-2

Trochet, A., Moulherat, S., Calvez, O., Stevens, V., Clobert, J., & Schmeller, D. (2014). A database of life-history traits of European amphibians. Biodiversity Data Journal, 2, e4123. https://doi.org/10.3897/BDJ.2.e4123

Van de Weyer, K., Meis, S., & Krautkrämer, V. (2015). Die Makrophyten des Großen Stechlinsees, des Wummsees und des Wittwesees. Fachbeiträge Des LUGV, 145, 1–92.

Van de Weyer, K., & Schmitt, C. (2011). Bestimmungsschlüssel für die aquatischen Makrophyten (Gefäßpflanzen, Armleuchteralgen und Moose) in Deutschland - Band 1: Bestimmungsschlüssel. Fachbeiträge Des LUGV, 119, 164pp.

Van der Maarel, E. (1979). Transformation of cover-abundance values in phytosociology and its effects on community similarity. Vegetatio, 39, 97–114. https://doi.org/10.1007/bf00052021

Venohr, M., Langhans, S. D., Peters, O., Hölker, F., Arlinghaus, R., Mitchell, L., & Wolter, C. (2018). The underestimated dynamics and impacts of water-based recreational activities on freshwater ecosystems. Environmental Reviews, 26, 199–213. https://doi.org/10.1139/er-2017-0024

Vilizzi, L., Tarkan, A. S., & Copp, G. H. (2015). Experimental evidence from causal criteria analysis for the effects of Common Carp (Cyprinus carpio) on freshwater ecosystems: A global perspective. Reviews in Fisheries Science and Aquaculture, 23, 253–290. https://doi.org/10.1080/23308249.2015.1051214

Wilson, A. M., Barr, J., & Zagorski, M. (2017). The feasibility of counting songbirds using unmanned aerial vehicles. The Auk, 134, 350–362. https://doi.org/10.1642/auk-16-216.1

Winfield, I. J. (2004). Fish in the littoral zone: Ecology, threats and management. Limnologica, 34, 124–131. https://doi.org/10.1016/S0075-9511(04)80031-8

Wright, S. W. (1991). Improved HPLC method for the analysis of chlorophylls and carotenoids from marine phytoplankton. Marine Ecology Progress Series, 77, 183–196. https://doi.org/10.3354/meps077183

Yoccoz, N. G., Nichols, J. D., & Boulinier, T. (2001). Monitoring of biological diversity in space and time. Trends in Ecology & Evolution, 16, 446–453. https://doi.org/10.1016/S0169-5347(01)02205-4

Zhao, T., Grenouillet, G., Pool, T., Tudesque, L., & Cucherousset, J. (2016). Environmental determinants of fish community structure in gravel pit lakes. Ecology of Freshwater Fish, 25, 412–421. https://doi.org/10.1111/eff.12222

