## supplementary information for "Status of aquatic and riparian biodiversity in artificial lake ecosystems with and without management for recreational fisheries: implications for conservation"

Table S1: Lake characteristics (morphology and trophic state). 16 top lakes = managed lakes (red), 10 bottom lakes = unmanaged lakes (green).

| **Lake name** | **Maximum  depth  in m (MaxDep)** | **Lake  area in  ha (LArea)** | **Shoreline development  factor (SDF)** | **Relative  depth  ratio (RelDepR)** | **Total phoshporus in µg*l^-1^ (TP)** | **Total organic carbon in mg*l^-1^ (TOC)** | **Mean chlorophyll a in µg*l^-1^**  **(CHLa)** | **Secchi depth  in m**  **(Secchi)** | **Ammonium in µg*l^-1^ (NH4)** | **Nitrate in µg*l^-1^ (NO3)** |
| --- | --- | --- | --- | --- | --- | --- | --- | --- | --- | --- |
| Chodhemster Kolk | 10.1 | 3.2 | 1.1 | 0.05 | 20 | 8.5 | 4.3 | 1.4 | 50 | 5 |
| Collrunge | 8.6 | 4.3 | 1.1 | 0.04 | 15 | 5.1 | 4.6 | 2.8 | 40 | 50 |
| Donner Kiesgrube 3 | 5.2 | 1.0 | 1.2 | 0.05 | 27 | 6.4 | 8.9 | 2.5 | 15 | 5 |
| Kiesteich Brelingen | 8.7 | 8.5 | 2.2 | 0.03 | 17 | 4.2 | 6.5 | 1.2 | 50 | 780 |
| Kolshorner Teich | 16.1 | 4.3 | 1.5 | 0.07 | 17 | 4.6 | 4.8 | 5.5 | 15 | 5 |
| Linner See | 11.2 | 17.7 | 1.8 | 0.02 | 18 | 6.6 | 7.4 | 4.0 | 15 | 5 |
| Meitzer See | 23.5 | 19.5 | 1.3 | 0.05 | 8 | 6.3 | 2.1 | 4.5 | 15 | 100 |
| Neumanns Kuhle | 6.2 | 6.9 | 1.1 | 0.02 | 160 | 13.0 | 65.3 | 0.5 | 15 | 5 |
| Plockhorst | 8.2 | 14.3 | 1.7 | 0.02 | 34 | 6.9 | 30.7 | 0.5 | 120 | 790 |
| Saalsdorf | 9.2 | 9.0 | 1.3 | 0.03 | 14 | 7.0 | 15.3 | 1.3 | 240 | 860 |
| Schleptruper See | 10.1 | 4.0 | 1.3 | 0.04 | 9 | 5.2 | 3.7 | 4.7 | 15 | 40 |
| Stedorfer Baggersee | 2.8 | 1.9 | 1.2 | 0.02 | 23 | 11.8 | 10.2 | 0.9 | 40 | 5 |
| Steinwedeler Teich | 9.1 | 10.4 | 2.0 | 0.03 | 10 | 2.5 | 5.8 | 3.0 | 40 | 120 |
| Wahle | 12.1 | 8.1 | 1.4 | 0.04 | 10 | 3.4 | 7.2 | 3.2 | 210 | 1040 |
| Weidekampsee | 4.3 | 2.9 | 1.6 | 0.02 | 10 | 5.4 | 3.0 | 2.8 | 15 | 5 |
| Wiesedermeer | 9.2 | 2.9 | 1.7 | 0.05 | 19 | 8.2 | 6.7 | 2.1 | 15 | 5 |
| Bülstedt | 1.1 | 2.4 | 1.6 | 0.01 | 25 | 6.7 | 90.6 | 1.6 | 30 | 2940 |
| Goldbeck | 5.0 | 2.3 | 1.4 | 0.03 | 19 | 2.9 | 19.2 | 2.3 | 70 | 770 |
| Handorf | 23.0 | 13.6 | 1.7 | 0.06 | 58 | 4.9 | 29.2 | 1.6 | 15 | 310 |
| Hänigsen | 12.3 | 6.2 | 1.4 | 0.04 | 16 | 5.5 | 9.8 | 2.7 | 50 | 2040 |
| Heeßel | 7.4 | 0.9 | 1.3 | 0.07 | 29 | 6.2 | 12.6 | 1.7 | 15 | 380 |
| Hopels | 14.5 | 5.5 | 1.3 | 0.05 | 12 | 5.6 | 4.8 | 4.5 | 50 | 140 |
| Lohmoor | 7.4 | 4.1 | 2.2 | 0.03 | 72 | 12.4 | 27.6 | 0.9 | 15 | 5 |
| Pfütze | 7.3 | 10.6 | 1.8 | 0.02 | 13 | 3.8 | 5.7 | 1.9 | 15 | 5 |
| Schwicheldt | 10.0 | 1.7 | 1.8 | 0.07 | 16 | 5.5 | 2.6 | 2.6 | 30 | 90 |
| Xella | 7.3 | 2.1 | 1.5 | 0.04 | 12 | 6.2 | 9.8 | 0.5 | 550 | 650 |
| **Mean ± SD** | **9.6 ± 5.2** | **6.5 ± 5.2** | **1.5 ± 0.3** | **0.04 ± 0.02** | **26.3 ± 30.9** | **6.3 ± 2.6** | **15.3 ± 20.0** | **2.4 ± 1.4** | **67.3 ± 111.8** | **428 ± 690** |

Table S2: Lake characteristics (trophic state and habitat structure). 16 top lakes = managed lakes (red), 10 bottom lakes = unmanaged lakes (green).

| **Lake name** | **Conductivity in µS*cm^-1^ (Con)** | **pH value (pH)** | **Volume-% of simple dead wood (SDW_Vol)** | **Volume-% of coarse woody structure (CWS_Vol)** | **Mean riparian tree coverage from 0 to 4 (Rip_Trees)** | **Mean litoral reed coverage from 0 to 4 (Reed)** | **Mean riparian vascular plant coverage**  **from 0 to 4 (Rip_Herbs)** | **Litoral submerged macrophyte coverage in % (MP_Cov)** | **Lake age in years by 2017 (Age)** |
| --- | --- | --- | --- | --- | --- | --- | --- | --- | --- |
| Chodhemster Kolk | 208 | 6.7 | 0.015 | < 0.1 | 0.4 | 0.2 | 3 | 22.4 | 46 |
| Collrunge | 200 | 7.9 | 0.003 | 0.4 | 0.8 | 2.4 | 1.3 | 67.9 | 35 |
| Donner Kiesgrube 3 | 632 | 7.8 | 0.002 | 2.1 | 1.5 | 0 | 1.7 | 38.5 | 17 |
| Kiesteich Brelingen | 299 | 7.5 | 0.006 | 1.4 | 0.8 | 0.1 | 1.7 | 12.5 | 18 |
| Kolshorner Teich | 547 | 7.6 | 0.001 | 1.1 | 1.0 | 1.3 | 1.8 | 37.9 | 37 |
| Linner See | 301 | 8.5 | 0.035 | 1.5 | 1.2 | 1.3 | 1.6 | 35.8 | 17 |
| Meitzer See | 618 | 7.9 | 0.003 | 0.8 | 0.9 | 2.5 | 1.8 | 33.6 | 11 |
| Neumanns Kuhle | 585 | 8.6 | 0.002 | 5.6 | 1.1 | 0.5 | 1.7 | 34.9 | 47 |
| Plockhorst | 338 | 8.0 | 0.002 | 1.0 | 1.2 | 1.5 | 1.5 | 27.6 | 19 |
| Saalsdorf | 566 | 9.0 | 0.004 | 1.2 | 0.8 | 2.3 | 1.6 | 19.5 | 22 |
| Schleptruper See | 502 | 8.2 | 0.004 | 2.0 | 1.1 | 2.0 | 1.7 | 17.4 | 52 |
| Stedorfer Baggersee | 352 | 7.5 | 0.001 | 0.3 | 1.2 | 0.3 | 2.1 | 59.8 | 34 |
| Steinwedeler Teich | 742 | 7.5 | 0.001 | 0.8 | 1.0 | 2.0 | 2.0 | 49.2 | 39 |
| Wahle | 728 | 7.6 | < 0.001 | 0.3 | 1.0 | 0.9 | 1.4 | 26.6 | 27 |
| Weidekampsee | 461 | 8.0 | < 0.001 | 3.2 | 1.0 | 2.2 | 1.9 | 82.3 | 23 |
| Wiesedermeer | 136 | 8.0 | 0.001 | 1.9 | 0.9 | 0.9 | 1.1 | 62.5 | 27 |
| Bülstedt | 285 | 7.6 | 0.023 | 1.6 | 1.2 | 1.7 | 0.2 | 7.6 | 26 |
| Goldbeck | 293 | 7.9 | 0.005 | 0.7 | 1.0 | 0.3 | 0.7 | 2.5 | 25 |
| Handorf | 666 | 9.2 | 0.002 | 0.2 | 0.6 | 0 | 0.1 | 8.3 | 13 |
| Hänigsen | 437 | 8.6 | 0.001 | 0.6 | 0.4 | 1.7 | 0.6 | 9.0 | 6 |
| Heeßel | 537 | 8.2 | 0.007 | 3.7 | 0.8 | 0 | 0.4 | 0 | 54 |
| Hopels | 215 | 7.7 | 0.028 | 6.2 | 1.2 | 0.8 | 0.9 | 8.2 | 19 |
| Lohmoor | 159 | 8.2 | 0.007 | 0.1 | 1.0 | 0.9 | 1.9 | 29.0 | 26 |
| Pfütze | 342 | 7.7 | 0.001 | 3.4 | 1.2 | 0.8 | 1.8 | 85.2 | 17 |
| Schwicheldt | 895 | 8.4 | 0.001 | < 0.1 | 0.5 | 0.7 | 1.7 | 52.9 | 10 |
| Xella | 957 | 7.5 | 0.001 | 0.9 | 1.2 | 1.7 | 1.8 | 8.5 | 42 |
| **Mean ± SD** | **462 ± 225** | **2.4 ± 1.4** | **0.006 ± 0.009** | **1.6 ± 1.6** | **1.0 ± 0.3** | **1.1 ± 0.8** | **1.5 ± 0.6** | **32.3 ± 23.8** | **27.3 ± 13.3** |

Table S3: Lake characteristics (land use, water and human presence). 16 top lakes = managed lakes (red), 10 bottom lakes = unmanaged lakes (green).

| **Lake name** | **Excavation in 100m-buffer in % (Excav)** | **Agriculture in 100m-buffer in % (Agric)** | **Forest in 100m-buffer in % (Forest)** | **Wetland in 100m-buffer in % (Wetland)** | **Water surface in 100m- buffer in % (Water)** | **Distance to next lake in m (DistLake)** | **Distance to next river in m (DistRiver)** | **Distance to next canal in m (DistCanal)** | **Distance to next road in m (DistRoad)** | **Urban area in 100m-buffer in % (Urban)** |
| --- | --- | --- | --- | --- | --- | --- | --- | --- | --- | --- |
| Chodhemster Kolk | 0 | 2.4 | 0 | 0 | 9.4 | 40 | 1170 | 1 | 135 | 87.5 |
| Collrunge | 0 | 34.7 | 0 | 0 | 1.6 | 170 | 29900 | 410 | 20 | 43.5 |
| Donner Kiesgrube 3 | 13.2 | 21.4 | 0 | 0 | 50.4 | 5 | 350 | 150 | 550 | 0 |
| Kiesteich Brelingen | 0 | 2.7 | 66.3 | 0 | 8.6 | 240 | 7540 | 1210 | 70 | 11.2 |
| Kolshorner Teich | 3.0 | 22.8 | 72.6 | 0 | 1.6 | 75 | 1850 | 350 | 630 | 0 |
| Linner See | 0 | 4.9 | 11.2 | 0 | 6.9 | 20 | 25 | 20 | 75 | 64.4 |
| Meitzer See | 0 | 4.3 | 68.4 | 0 | 15.0 | 40 | 425 | 1630 | 50 | 8.0 |
| Neumanns Kuhle | 0 | 3.2 | 8.4 | 14.4 | 0.9 | 850 | 4800 | 1 | 40 | 55.1 |
| Plockhorst | 0 | 14.5 | 17.2 | 0 | 19.2 | 40 | 1130 | 275 | 15 | 30.6 |
| Saalsdorf | 0 | 35.5 | 5.9 | 0 | 2.7 | 80 | 1470 | 30 | 520 | 55.9 |
| Schleptruper See | 0 | 8.3 | 24.3 | 0 | 2.0 | 180 | 500 | 1 | 40 | 32.6 |
| Stedorfer Baggersee | 0 | 6.5 | 11.9 | 0 | 2.2 | 5 | 1320 | 30 | 1010 | 56.1 |
| Steinwedeler Teich | 11.6 | 55.9 | 25.9 | 0 | 1.7 | 185 | 38 | 360 | 20 | 0 |
| Wahle | 15.3 | 44.0 | 9.1 | 0 | 14.4 | 30 | 710 | 250 | 35 | 0 |
| Weidekampsee | 21.3 | 47.7 | 1.7 | 0 | 15.9 | 55 | 2600 | 30 | 365 | 0 |
| Wiesedermeer | 11.7 | 55.8 | 29.7 | 0 | 2.9 | 610 | 29790 | 250 | 670 | 0 |
| Bülstedt | 0 | 52.0 | 0 | 4.3 | 8.0 | 40 | 820 | 1180 | 640 | 0 |
| Goldbeck | 14.0 | 6.6 | 5.6 | 0 | 13.1 | 15 | 555 | 110 | 150 | 51 |
| Handorf | 1.1 | 20.2 | 0 | 0 | 6.1 | 30 | 770 | 465 | 230 | 59.5 |
| Hänigsen | 32.2 | 18.3 | 5.5 | 0 | 0.5 | 1280 | 560 | 305 | 35 | 43.5 |
| Heeßel | 1.3 | 10.2 | 15.4 | 0 | 14.0 | 10 | 820 | 65 | 30 | 0 |
| Hopels | 6.5 | 79.0 | 0 | 1.6 | 2.1 | 330 | 31920 | 5 | 1530 | 0 |
| Lohmoor | 0 | 51.9 | 0 | 45.1 | 1.7 | 830 | 220 | 15 | 1150 | 0 |
| Pfütze | 0 | 54.0 | 8.0 | 0 | 30.1 | 45 | 480 | 40 | 445 | 0 |
| Schwicheldt | 39.0 | 42.0 | 15.5 | 0 | 3.6 | 60 | 3050 | 15 | 975 | 0 |
| Xella | 0 | 3.5 | 6.3 | 0 | 8.0 | 1 | 800 | 45 | 395 | 20.1 |
| **Mean ± SD** | **6.5 ± 10.4** | **27.0 ± 22.4** | **15.7 ± 21.4** | **2.5 ± 9.2** | **9.3 ± 11.0** | **202.5 ± 325.4** | **4754.3 ± 9639.7** | **278.6 ± 422.9** | **377.9 ± 417.2** | **23.8 ± 27.5** |

Table S4: Lake characteristics (human presence and recreational intensity). 16 top lakes = managed lakes (red), 10 bottom lakes = unmanaged lakes (green).

| **Lake name** | **Distance to next village or city in m (DistVille)** | **Distance to next city in m (DistCity)** | **Litter related to angling in No.*m_shore_^-1^ (A_Lit)** | **Litter not related to angling in No.*m_shore_^-1^  (NonA_Lit)** | **Angling-sites and open spaces in % of shoreline (open_sites)** | **Trails and paths per shoreline in m*m^-1^ (Trails)** | **Anglers per visit (Anglers)** | **Dog walker and dogs per visit (Dogs)** | **swimming people  per visit (Swimmers)** | **other people per visit (other_ people)** |
| --- | --- | --- | --- | --- | --- | --- | --- | --- | --- | --- |
| Chodhemster Kolk | 170 | 170 | 0.02 | 0.1 | 87.7 | 1.0 | 0 | 3.6 | 2.2 | 6.6 |
| Collrunge | 20 | 9000 | 0 | < 0.1 | 6.3 | 0.8 | 0.5 | 0.5 | 0.7 | 0.8 |
| Donner Kiesgrube 3 | 600 | 3650 | 0 | 0.3 | 3.6 | 1.0 | 0 | 0 | 0 | 0.3 |
| Kiesteich Brelingen | 150 | 10500 | 0.03 | 1.0 | 17.2 | 0.8 | 1.3 | 5.2 | 4.2 | 11.9 |
| Kolshorner Teich | 460 | 4180 | 0.19 | 1.0 | 16.4 | 1.0 | 4.2 | 0 | 4.4 | 1.4 |
| Linner See | 380 | 7630 | 0.10 | 1.5 | 14.3 | 1.0 | 5.1 | 0.6 | 6.0 | 0.6 |
| Meitzer See | 1160 | 8580 | 0.06 | 0.8 | 8.5 | 1.0 | 3.1 | 6.0 | 10 | 5.5 |
| Neumanns Kuhle | 1400 | 7470 | 0.01 | 0.5 | 5.5 | 1.0 | 0.6 | 0.4 | 0.7 | 0.6 |
| Plockhorst | 180 | 11180 | 0.04 | 1.2 | 16.4 | 1.0 | 1.0 | 2.4 | 2.4 | 5.7 |
| Saalsdorf | 825 | 13060 | 0.06 | 0.6 | 18.0 | 1.0 | 1.2 | 0.6 | 1.9 | 1.0 |
| Schleptruper See | 200 | 1540 | 0.02 | 0.3 | 17.2 | 1.1 | 1.7 | 1.6 | 2.5 | 1.9 |
| Stedorfer Baggersee | 240 | 5240 | 0.04 | 0.4 | 27.4 | 1.1 | 0.5 | 0.4 | 0.7 | 0.4 |
| Steinwedeler Teich | 775 | 1230 | 0.13 | 1.5 | 17.1 | 0.8 | 4.2 | 1.6 | 5.3 | 3.0 |
| Wahle | 320 | 8000 | 0 | 0.4 | 25.8 | 1.0 | 1.2 | 3.5 | 3.0 | 4.5 |
| Weidekampsee | 225 | 13130 | 0.01 | 1.5 | 6.1 | 0.8 | 0.3 | 0.5 | 0.6 | 1.7 |
| Wiesedermeer | 960 | 9600 | 0.04 | 0.2 | 8.1 | 0.6 | 0.8 | 0.7 | 1.4 | 0.4 |
| Bülstedt | 240 | 11700 | 0 | 0 | 4.9 | 0.2 | 0.3 | 0 | 0.3 | 0.3 |
| Goldbeck | 300 | 5960 | 0.02 | < 0.1 | 7.8 | 0.2 | 0 | 0 | 0.8 | 0.5 |
| Handorf | 150 | 1070 | 0 | 0.1 | 4.6 | 0.9 | 0 | 3.3 | 3.1 | 3.4 |
| Hänigsen | 60 | 4880 | 0 | 0.5 | 4.6 | 1.4 | 0.8 | 0 | 1.0 | 3.8 |
| Heeßel | 710 | 2080 | 0 | 0 | 7.0 | 0.2 | 0 | 0 | 0 | 0 |
| Hopels | 1400 | 15110 | 0 | 2.3 | 0.8 | 0.6 | 0 | 0.7 | 0.4 | 0.4 |
| Lohmoor | 1810 | 7180 | 0 | < 0.1 | 1.1 | 0.1 | 0 | 0 | 0 | 0.1 |
| Pfütze | 440 | 5300 | 0 | 0.4 | 6.0 | 0.5 | 0 | 0.9 | 1.1 | 0.7 |
| Schwicheldt | 1150 | 2010 | 0 | 0 | 0 | 0 | 0 | 0 | 0 | 0 |
| Xella | 1750 | 3300 | 0 | 0.1 | 3.0 | 0 | 0.2 | 0 | 0.2 | 0 |
| **Mean ± SD** | **618.3 ± 533.4** | **6644.2 ± 4204.5** | **0.03 ± 0.05** | **0.6 ± 0.6** | **14.6 ± 17.9** | **0.7 ± 0.4** | **1.0 ± 1.5** | **1.2 ± 1.7** | **2.0 ± 2.4** | **2.1 ± 2.8** |

Table S5: PCA-axes and their interpretation of classes of environmental variables

Only the first four axes are shown. Other axes had eigenvalues < 1.

| **Characteristic** | ***Variable*** | **Dim 1** | **Dim 2** | **Dim 3** | **Dim 4** |
| --- | --- | --- | --- | --- | --- |
| **Morphology** | *Eigenvalue* | 1.71 | 1.47 | 0.75 | 0.07 |
|  | *Proportion of explained variance in %* | 43 | 37 | 19 | 2 |
|  | *MaxDep* | -0.75 | 0.09 | -0.03 | -0.66 |
|  | *LArea* | -0.54 | -0.51 | -0.39 | 0.56 |
|  | *SDF* | -0.07 | -0.59 | 0.81 | -0.04 |
|  | *RelDepR* | -0.39 | 0.63 | 0.45 | 0.51 |
|  | *Interpretation of axes* | lake shallowness | lake steepness | *(not used)* | *(not used)* |
| **Trophic state** | *Eigenvalue* | 2.65 | 1.69 | 1.32 | 1.15 |
|  | *Proportion of explained variance in %* | 33 | 21 | 16 | 14 |
|  | *TP* | 0.53 | -0.14 | 0.27 | -0.06 |
|  | *TOC* | 0.47 | -0.23 | 0.13 | 0.33 |
|  | *CHLa* | 0.50 | 0.18 | -0.29 | -0.24 |
|  | *Secchi* | -0.43 | -0.26 | -0.01 | -0.40 |
|  | *NH4* | -0.05 | 0.62 | 0.20 | 0.37 |
|  | *NO3* | 0.13 | 0.49 | -0.54 | -0.32 |
|  | *Con* | -0.11 | 0.44 | 0.58 | -0.15 |
|  | *pH* | 0.21 | 0.00 | 0.40 | -0.64 |
|  | *Interpretation of axes* | trophic state | nitrogen | conductivity | acidity |
| **Habitat structure** | *Eigenvalue* | 1.76 | 1.50 | 0.95 | 0.80 |
|  | *Proportion of explained variance in %* | 29 | 25 | 16 | 13 |
|  | *SDW_Vol* | -0.46 | 0.24 | -0.37 | 0.62 |
|  | *CWS_Vol* | -0.33 | 0.56 | 0.18 | -0.31 |
|  | *Rip_Trees* | -0.04 | 0.68 | -0.04 | -0.08 |
|  | *Reed* | 0.32 | 0.09 | -0.88 | -0.24 |
|  | *Rip_Herbs* | 0.51 | 0.22 | 0.14 | 0.66 |
|  | *MP_Cov* | 0.56 | 0.34 | 0.18 | -0.14 |
|  | *Interpretation of axes* | vegetated habitat | woody habitat | *(not used)* | *(not used)* |

Table S6: PCA-axes and their interpretation for environmental variables.

Only the first four axes are shown. Other axes had eigenvalues < 1.

| **Characteristic** | ***Variable*** | **Dim 1** | **Dim 2** | **Dim 3** | **Dim 4** |
| --- | --- | --- | --- | --- | --- |
| **Land use  (in 100 m   around lake)** | *Eigenvalue* | 1.77 | 1.23 | 0.79 | 0.22 |
|  | *Proportion of explained variance in %* | 44 | 31 | 20 | 5 |
|  | *Excav* | 0.44 | -0.23 | -0.86 | -0.08 |
|  | *Agric* | 0.65 | -0.17 | 0.43 | -0.61 |
|  | *Forest* | -0.06 | 0.86 | -0.21 | -0.46 |
|  | *Urban* | -0.62 | -0.42 | -0.15 | -0.65 |
|  | *Interpretation of axes* | agricultural extent | forest extent | *(not used)* | *(not used)* |
| **Water** | *Eigenvalue* | 1.82 | 1.15 | 0.96 | 0.65 |
|  | *Proportion of explained variance in %* | 36 | 23 | 19 | 13 |
|  | *Wetland* | 0.50 | -0.52 | 0.23 | 0.24 |
|  | *Water* | -0.51 | -0.33 | -0.05 | 0.78 |
|  | *DistLake* | 0.63 | -0.08 | 0.15 | 0.24 |
|  | *DistRiver* | 0.28 | 0.71 | -0.29 | 0.51 |
|  | *DistCanal* | -0.17 | 0.34 | 0.92 | 0.11 |
|  | *Interpretation of axes* | wetland | distance to next river | *(not used)* | *(not used)* |
| **Human presence** | *Eigenvalue* | 1.57 | 0.95 | 0.48 | - |
|  | *Proportion of explained variance in %* | 52 | 32 | 16 | - |
|  | *DistRoad* | 0.69 | -0.06 | -0.72 | - |
|  | *DistVille* | 0.62 | -0.47 | 0.63 | - |
|  | *DistCity* | 0.38 | 0.88 | 0.28 | - |
|  | *Interpretation of axes* | rural | *(not used)* | *(not used)* | *(no 4^th^ dimension)* |
| **Recreational use** | *Eigenvalue* | 3.61 | 1.81 | 0.93 | 0.79 |
|  | *Proportion of explained variance in %* | 45 | 23 | 12 | 10 |
|  | *A_Lit* | 0.36 | 0.41 | -0.35 | 0.10 |
|  | *NonA_Lit* | 0.29 | 0.28 | 0.42 | -0.46 |
|  | *open_sites* | 0.14 | -0.39 | -0.76 | -0.23 |
|  | *Trails* | 0.34 | -0.06 | -0.01 | -0.69 |
|  | *Anglers* | 0.42 | 0.38 | -0.17 | 0.19 |
|  | *Dogs* | 0.36 | -0.47 | 0.24 | 0.26 |
|  | *Swimmers* | 0.48 | 0.03 | 0.06 | 0.39 |
|  | *other_people* | 0.35 | -0.48 | 0.20 | 0.00 |
|  | *Interpretation of axes* | general recreational use intensity | inaccessibility | *(not used)* | *(not used)* |

Table S7: Variance inflation factors (VIFs) for explanatory variables (axes from Tables S5, S6).
 VIFs > 5 indicate highly correlated variables in the multivariate space.

| ***Variable*** | **VIF** |
| --- | --- |
| lake shallowness | 4.68 |
| lake steepness | 2.43 |
| vegetated habitat | 2.50 |
| woody habitat | 3.45 |
| trophic state | 3.84 |
| nitrogen | 1.81 |
| conductivity | 3.07 |
| acidity | 3.64 |
| agricultural extent | 3.24 |
| forest extent | 1.82 |
| wetland | 2.94 |
| distance to next river | 1.54 |
| rural | 4.98 |
| general recreational use intensity | 4.36 |
| inaccessibility | 2.82 |
| age | 3.06 |

Table S8: PCA-axes and their interpretation for species richness including the loadings of explanatory variables from RDA. Names of these variables are interpretations from Table S5, S6. Only the first four axes are shown. Other axes had eigenvalues < 1.

| ***Species richness*** | **Dim 1** | **Dim 2** | **Dim 3** | **Dim 4** |
| --- | --- | --- | --- | --- |
| *Eigenvalue* | *2,19* | *1,56* | *1,28* | *0,95* |
| *Proportion of variance explained in %* | *31,27* | *22,24* | *18,28* | *13,63* |
| submerged macrophytes | 0,20 | 0,78 | 0,48 | -0,14 |
| riparian_vascular plants | 0,56 | -0,60 | 0,41 | -0,24 |
| riparian_trees | -0,48 | 0,25 | 0,78 | -0,02 |
| amphibians | 0,75 | 0,21 | 0,06 | -0,21 |
| Odonata | 0,64 | 0,33 | -0,31 | -0,42 |
| songbirds | 0,65 | -0,43 | 0,40 | 0,38 |
| waterfowl | 0,48 | 0,44 | -0,13 | 0,71 |
| *Interpretation of Axes* | *Riparian Species Richness* | *Submerged Macrophytes* | *Riparian Tree Species Richness* | *Waterfowl Species Richness* |
| woody_habitat | -0,57 | 0,37 | 0,26 | -0,05 |
| agricultural_extent | 0,44 | 0,26 | 0,19 | -0,30 |
| lake_steepness | -0,21 | -0,28 | -0,33 | -0,41 |

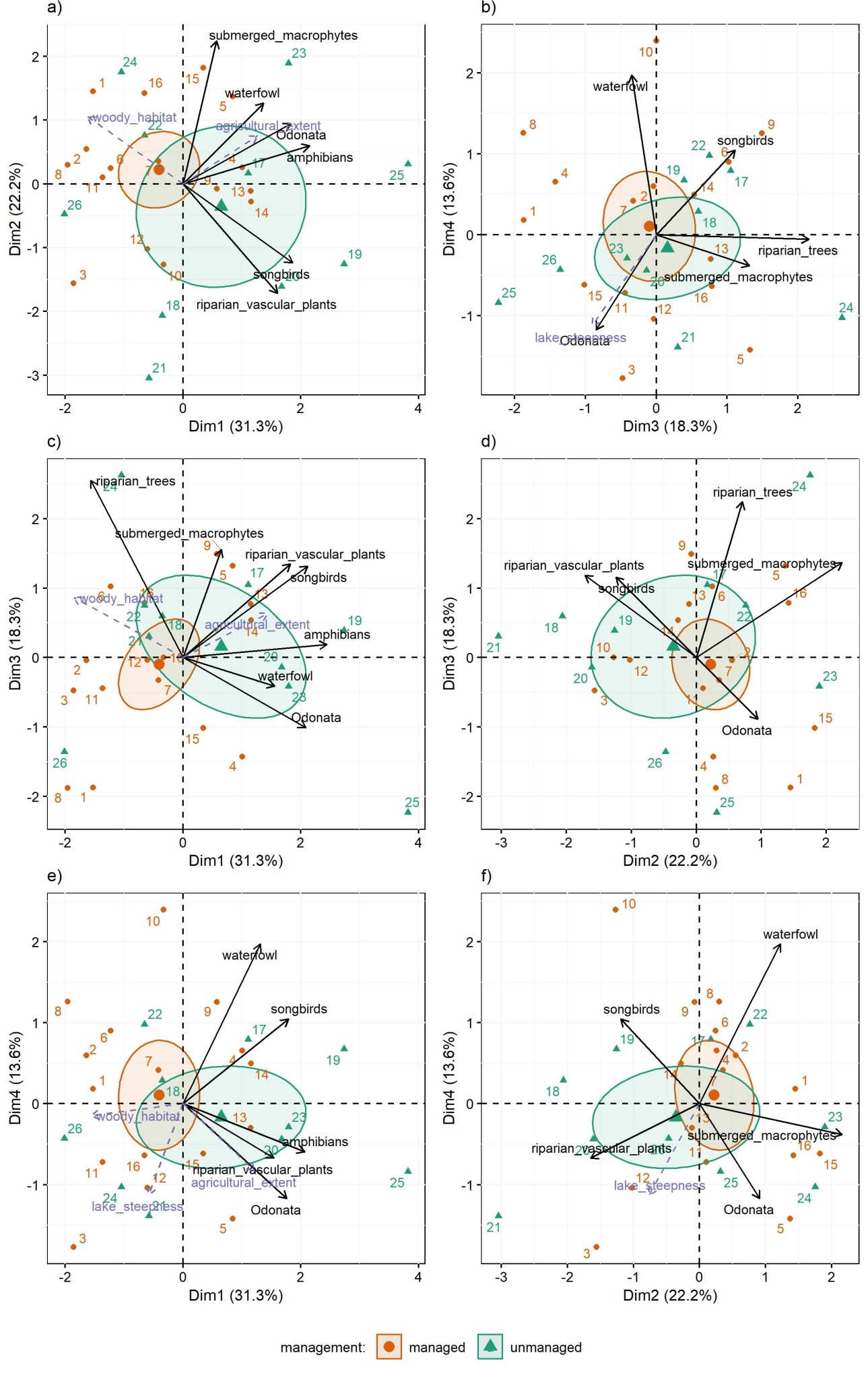

Figure S1: Principal component analysis (PCA) of species richness plotted for the first 4 axes (a: Dim 1 and 2, b: Dim 3 and 4, c: Dim 1 and 3, d: Dim 2 and 3, e: Dim 1 and 4, f: Dim 2 and 4). Percentages in brackets show the proportional variance explained by each axis. Only variables contributing to each plot are shown respectively. Numbers reflect the different lakes (Table 1). The centroids of management types and the explanatory variables from redundancy analysis (RDA, slashed purple lines) are plotted as supplementary variables to not influence the ordination. The 95% confidence-level around centroids are plotted to visualize differences between lake types.
